# Specific non-myogenic mesenchymal cells contribute to rotator cuff tear myosteatosis and fibrosis revealing novel therapeutic options

**DOI:** 10.64898/2026.02.11.705391

**Authors:** Helen Rueckert, Anthony J. Mirando, Abigail P. Leinroth, Juliana Ibarra, Joe V. Chakkalakal, Matthew J. Hilton

## Abstract

The rotator cuff is a group of four muscles in the shoulder, which aid in movement and rotation of the upper arm. Rotator cuff tears (RCTs) within tendons of these muscles are common musculoskeletal injuries, often resulting in intramuscular fat, fibrosis, and muscle atrophy. Fatty infiltration specifically correlates with high rates of retear following repair. The cellular sources and molecular cues that cause these pathologies are unknown and therefore non-surgical cell/drug therapies for RCTs do not exist. Thus, we first sought to determine the cellular source(s) and molecular underpinnings of fatty atrophy and fibrosis associated with massive RCTs. Using a murine model of massive RCTs combined with lineage tracing, we demonstrate that muscle resident *Pdgfra+* non-myogenic mesenchymal cells (NMMCs) are responsible for the fatty and fibrotic RCT pathologies. Utilizing sorted *Pdgfra+* cells from rotator cuff muscles and “deep” single cell RNA-sequencing, we identified a specific *Dpp4+* cell population associated with RCT-induced fibrosis, while *Gfra1+* nerve-associated NMMCs are drivers of the RCT-induced intramuscular fat pathology. Finally, we demonstrate that RCT-induced fatty infiltration occurs at least partially via the loss of GDNF-GFRA1-RET signaling, since local treatment of murine RCTs with a small molecule RET agonist reduces development of the RCT-induced intramuscular fat.

## INTRODUCTION

The rotator cuff is a group of four muscles (supraspinatus, infraspinatus, subscapularis, and teres minor) in the shoulder, which aid in movement and rotation of the upper arm. Rotator cuff tears (RCTs) within tendons of these muscles are some of the most common shoulder ailments and can result in intramuscular fat development (known as fatty atrophy or infiltration), fibrosis, and muscle atrophy (1–3). RCTs vary in size, from partial tears to massive, multi-muscle, full thickness tears. RCTs are clinically diagnosed via MRI and depending on size can be treated with physical therapy, repair surgery, or full shoulder arthroplasty. RCT repairs are extremely common in the US with over 275,000 surgeries occurring each year (4). Unfortunately, the rates of retear following repair are significant—as high as 94% for certain tear types/sizes (5)—and clinical work has correlated this to high levels of MRI diagnosed intramuscular fatty infiltration (6–8). This MRI Goutallier classification of intramuscular fat has also been confirmed histologically in RCT patient muscle biopsies that show significantly higher intramuscular lipid levels correlate with increasing size (small to massive) of full tears (9, 10) Unfortunately, the cellular sources and molecular cues that cause the pathologies of fat, fibrosis, and atrophy remain unknown and therefore there are no therapeutic treatment options for these RCT-induced pathologies, especially fatty atrophy (11–13). There is evidence that tendon injury in the rotator cuff results in more intramuscular fat than other muscle groups such as the gastrocnemius (Achilles tendon) or biceps brachii (bicep tendon) (14), suggesting the rotator cuff muscle group may have unique properties or unique cell populations that contribute to these pathologies. Muscle stem cells, *Pdgfra+* non-myogenic mesenchymal cells (NMMCs), and extra-muscular pre-adipocytes have all been hypothesized to contribute to these fibrotic and fatty atrophy pathologies (15–18).

Pdgfra+ NMMCs, sometimes referred to as fibro-adipogenic progenitor cells (FAPs), are a population of mesenchymal cells in the skeletal muscle niche first described in 2010 by Joe et al. and Uezumi et al. (19, 20). They are muscle niche cells defined by their expression of *Pdgfra* and *Sca1* in murine systems and solely *Pdgfra* in humans and other animal models (19–22). These NMMCs are found interstitial to the muscle surrounding muscle fibers, and are localized circumferentially around vessels, nerves and at neuromuscular junctions within skeletal muscle. Early work showed the ability of these Pdgfra+ cells to differentiate into fibroblasts, adipocytes, and chondrocytes (19, 23), as well as an ability to undergo population expansion following skeletal muscle injury. Subsequent studies evaluated these cells in regenerative muscle injury contexts, defining their expansion peaking at 3-5 days-post-injury (dpi) and demonstrating roles in transient ECM remodeling, signaling to a variety of immune cells and supporting activated satellite cells, before undergoing programmed cell death to return to homeostatic population levels during skeletal muscle regeneration (19, 24–28). Further work found that the ablation of Pdgfra+ cells dramatically hinders muscle regeneration (29) and negatively impacts muscle health during development and aging (29, 30). During development, TCF7L2+ progenitors that give rise to Pdgfra+ cells are required for proper skeletal muscle patterning and growth (31, 32). Several studies have implicated this broad classification of mesenchymal cells with a number of skeletal muscle pathologies where their aberrant proliferation and differentiation leads to intramuscular fibrosis, adipogenesis, and even ossification (well-reviewed in (28, 33–35)).

In recent years, cellular heterogeneity has gained interest in NMMC/FAP biology. Single cell RNA-sequencing (scRNA-seq) and single nucleus RNA-sequencing (snRNA-seq) studies have identified a few subpopulations of NMMCs/FAPs present during skeletal muscle development, following muscle injury, and in certain disease contexts (27, 36–40). These studies have largely performed unbiased scRNA-seq/snRNA-seq in which NMMCs/FAPs are just a fraction of the cells isolated and sequenced from skeletal muscle, which often limits the depth of sequencing (typically 15,000-30,000 reads per cell) and identifies “cell types” based on a limited number of highly expressed genes. Recently, we performed “deep” scRNA-seq (∼60,000 reads per cell) of NMMCs isolated from juvenile hindlimb skeletal muscles and identified novel NMMC subpopulations with some demonstrating unique differentiation potentials, response to injuries, or localization within hindlimb skeletal muscles (37). This study was the first of its kind to further define NMMCs/FAPs at the functional and spatial level in addition to their refined identification using “deep” scRNA-seq approaches. Here we have expanded upon these findings through the “deeper” sequencing (120,000-165,000 reads per cell) of NMMCs derived from forelimb rotator cuff muscles (supraspinatus and infraspinatus) in both uninjured and RCT conditions. Since RCT-induced muscle pathologies share common phenotypes (intramuscular fat and fibrosis) associated with several skeletal muscle diseases often attributed to aberrant regulation and/or differentiation of NMMCs/FAPs, we set out to explore the heterogeneity of Pdgfra*+* NMMCs in the rotator cuff muscles (forelimb muscles as opposed to hindlimb muscles), assess potential subpopulation specific cellular contributions to the murine massive RCT pathologies, and establish our “deeper” scRNA-sequencing approach as a method for not only identifying potentially unique NMMC populations, but also uncovering some of the molecular mechanisms driving these pathologies.

## RESULTS

### Murine Model of Massive Rotator Cuff Tears

To study massive RCT pathologies at the whole muscle and cellular level it was necessary to utilize an animal model of massive RCTs. Multiple groups have developed models of full tendon tears with or without nerve injury, and even combining these surgeries with complete removal of humoral head to prevent any tendon reattachment (41–43). Among these murine models it was determined that full tendon tears to the supraspinatus (SS) and infraspinatus (IS) coupled with damage to the suprascapular nerve (SSN) produced the most clinically relevant levels of fatty infiltration, fibrosis, and muscle atrophy (***Fig. 1A***). Though under-explored clinically, nerve involvement/damage is suspected to occur in many RCTs given the location of the SSN tract. Cadaver studies indicate that following massive RCTs, the supra- and infraspinatus muscle retract resulting in pinching or damage to the SSN at both the suprascapular notch and/or the spinoglenoid fossa (44). SSN neuropathies diagnosed by altered EMG and reduced nerve conduction have been reported following rotator cuff damage (45, 46). We and others have demonstrated that tendon injury alone in mice does not recapitulate the levels of fat generation observed in massive human RCTs, consistent with nerve involvement in the human pathology. Here we first assessed the levels of fat, fibrosis, and atrophy in this (SS+IS+SSN) model through whole muscle qPCR. At 8 weeks post-injury (8wpi), adipogenic genes (*Pparg* and *Plin1*), fibrotic markers (*Col1a1* and *Fn1*), and skeletal muscle atrophy genes (*Fbox32* and *Trim63)* were all significantly elevated in RCT muscles as compared to controls (***Fig. 1B*)**. Further quantification of Oil red O (fat stain) and FIBRONECTIN percent area (fibrosis) show significant increases in levels of these pathologies (***Fig. 1C,D***). Histologic stains (H&E and picrosirius red) show disruption in the myofibers with an increase in interstitial adipocytes (***Fig. 1E***) as well as increases in fibrosis (***Fig. 1F***). This in-depth model characterization demonstrates that the murine massive RCT injury model of SS+IS+SSN damage recapitulates human RCT muscle associated phenotypes.

**Figure 1:**
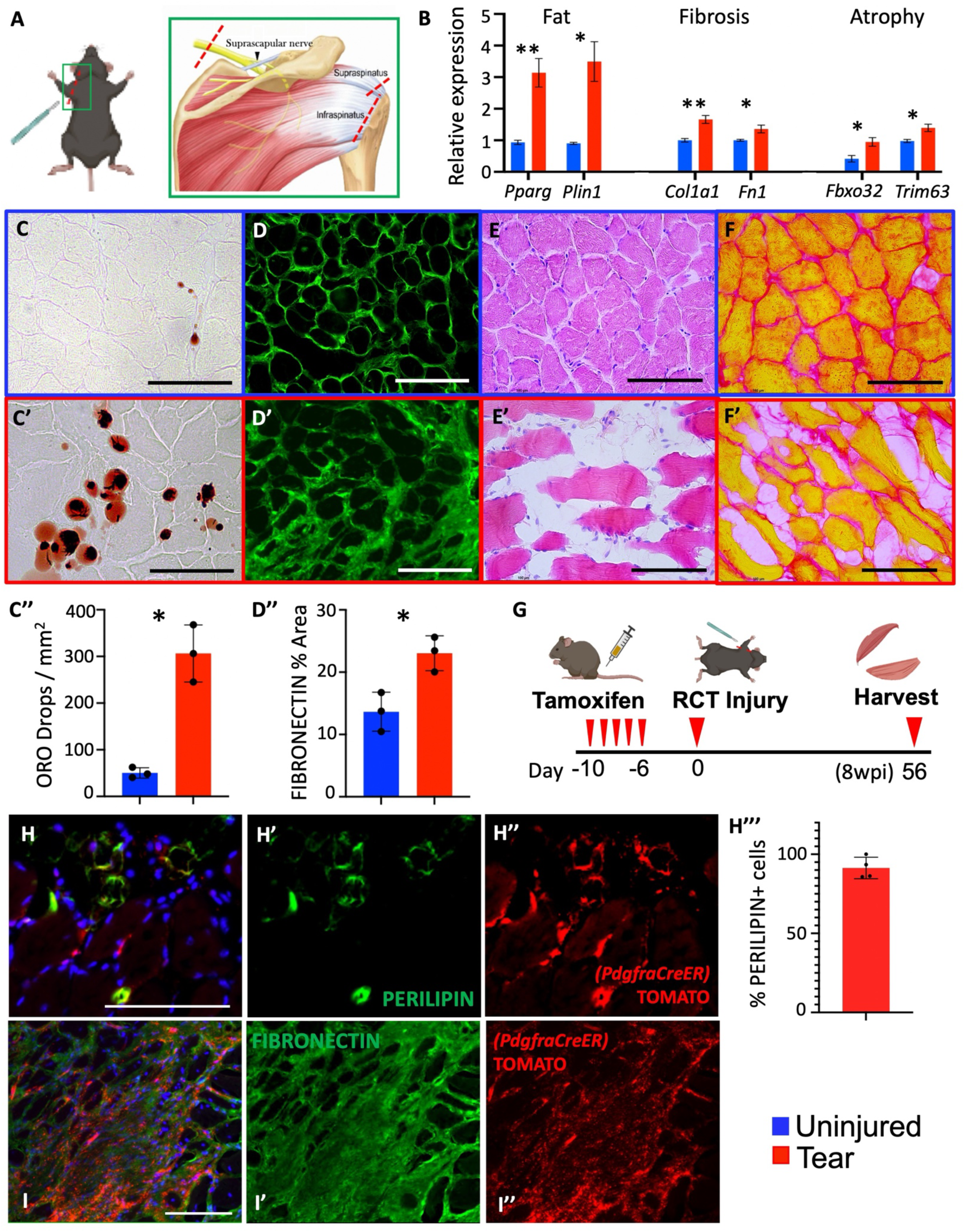
Massive RCT pathologies are caused by *Pdgfra*+ NMMCs. A) diagram of massive RCT injury showing areas of surgical damage on supraspinatus tendon, infraspinatus tendon, and suprascapular nerve. B) whole muscle qPCR from supraspinatus muscles 8wpi massive RCT. Data are represented as mean±SEM; *Pparg* p=0.009, *Plin1* p=0.015, *Col1a1* p=0.008, *Fn1* p=0.048, *Fbxo32* (ATROGIN1) p=0.037, *Trim63* (MURF1) p=0.030; N=3. Oil Red O staining for contralateral control (C) and 8wpi tear supraspinatus (C’) muscles with quantification of differences (C’’). Data are represented as mean±SD, p=0.0149; n=3, N=3. Significance determined using paired t-test. D) Fibronectin staining for contralateral control (D) and 8wpi tear supraspinatus (D’) muscles with quantification of differences (D’’). Data are represented as mean±SD, p=0.0242; n=3, N=3. Significance determined using paired t-test. E) Hemotoxylin and Eosin (H&E) stain of contralateral control (E) and 8wpi tear supraspinatus (E’) muscles. F) Picrosirus red stain of contralateral control (F) and 8wpi tear supraspinatus (F’) muscles. G) Schematic of tamoxifen injection and injury timeline for lineage tracing experiment. H) Representative image of colocalization of PERILIPIN (H’) and TOMATO (H’’) with quantification(H’’’) of PERILIPIN+ cells which are also TOMATO+ from the 8wpi *PdgfraCreER;R26-Ai9^fx/fx^* supraspinatus; n=4, N=3 I) Representative image of colocalization of FIBRONECTIN (H’) and TOMATO (I’’) from the 8wpi *PdgfraCreER;R26-Ai9^fx/fx^* supraspinatus; n=4, N=3. All scale bars = 100um. Schematics crafted with Biorender.

### Lineage Tracing of *Pdgfra+* NMMCs

After establishing this massive RCT model in the mouse, we sought to perform lineage tracing studies to determine the cellular source of RCT induced intramuscular fat and fibrosis. Given their known differentiation capacities, muscle resident *Pdgfra+* NMMCs appeared to be a likely source. Additionally, RCTs performed on *Pdgfra-H2b-Gfp*+ mice, have identified GFP+/PDGFRA+ cells near sites of intramuscular fat and fibrosis (17, 47). To determine whether these cells truly give rise to adipocytes and fibrotic fibroblasts, we performed lineage tracing using adult *PdgfraCre^ER^;R26-Ai9^fx/fx^* mice injected with tamoxifen and then subjected to massive RCT injury followed by rotator cuff muscle isolation at 8wpi (***Fig. 1G***). Demonstrating their non-myogenic nature, tdTOMATO+ myofibers are never observed following tamoxifen administration in uninjured or RCT injured muscles (data not shown). Immunofluorescent staining demonstrates that 91.3% of PERILIPIN+ mature adipocytes colocalized with tdTOMATO signal (***Fig. 1H***), indicating that nearly all RCT-induced adipocytes were derived from *Pdgfra+* NMMCs. Additionally, large areas of tdTOMATO positivity colocalized with plaques of the extracellular matrix (ECM) protein FIBRONECTIN (***Fig. 1I***), suggesting *Pdgfra+* cells were primarily responsible for the secretion of an excessive and fibrotic ECM. Thus, *Pdgfra+* NMMCs are the muscle niche cell population responsible for the RCT induced pathologies of fatty infiltration and fibrosis.

### Single Cell RNA Sequencing of *Pdgfra+* NMMCs

When first discovered, muscle resident NMMCs were treated as a homogeneous *Pdgfra*-expressing mesenchymal cell population; however, several recent scRNA-seq/snRNA-seq studies have found cellular heterogeneity and identified subpopulations within the *Pdgfra+* cells. To date, no single cell sequencing work has been performed on the rotator cuff muscles (whole muscle or NMMC specific)—so we sought to investigate this in uninjured and post-RCT-associated muscles. First, we assessed NMMC population dynamics following RCT injury in the mouse. Using *Pdgfra-H2b-Gfp* reporter mice, we performed massive RCT injuries and harvested RCT muscles at 5dpi, 2wpi, and 4wpi to identify *Pdgfra+* cell population expansion and contraction dynamics. Using fluorescent activated cell sorting (FACS) of live GFP+ cells from total SS and IS muscles, we found that the number of GFP+ cells were dramatically increased in RCT muscles at 5dpi compared to the uninjured contralateral rotator cuff muscles (65.1%), and at 2wpi they remained elevated (19.4%) (***Supp. Fig. 1A***). However, by 4wpi the levels of Pdgfra+/GFP+ cells had decreased compared to the controls (−20.2%) (***Supp. Fig. 1A***). Additionally, histologic analyses at 2wpi indicate slight increases in ORO and FIBRONECTIN levels that were lower than the 8wpi timepoint, suggesting that the 2wpi timepoint marks a period of NMMC expansion that is just beginning the pathogenic differentiation process contributing to fat and fibrosis (***Supp. Fig. 1B,C***). Since we sought to understand the NMMCs following peak expansion and as they initiate a pathogenic differentiation trajectory, we determined 2wpi as the most appropriate post-injury analysis timepoint.

To isolate Pdgfra+ NMMCs, we utilized the adult *Pdgfra-H2b-Gfp* reporter mouse to perform scRNA-seq on uninjured and 2wpi rotator cuff sorted GFP+ NMMCs to capture all NMMCs following injury activation and prior to their overt differentiation. SS and IS muscles were harvested, mechanically and enzymatically digested, live GFP+ cells were subjected to FACS, then single cell RNA libraries were prepared and sequenced (***Fig. 2A*)**. Our deep sequencing of sorted single cells generated 119,688 reads/cell in uninjured muscles and 164,517 reads/cell in RCT muscles leading to a high median number of genes identified per cell (4,517 and 5328, respectively). Following standard pre-processing, sequencing data from 2556 cells from uninjured muscles and 1994 cells from RCT muscles at 2wpi were integrated using the Seurat Platform (48) and projected as a UMAP, showing 9 distinct clusters of *Pdgfra+* cells (***Fig. 2B***). Control and 2wpi plots were also separately visualized (***Fig. 2C***) and fold-change subpopulation analyses showed increased cell numbers in some clusters (2, 7, 9), while others decreased (3, 5, 6, 8) or remained unchanged (1, 4) (***Fig. 2D***). Some previously defined NMMC subpopulation marker genes as well as novel rotator cuff specific genes were used to assess the identities of specific clusters. Clusters 1, 3, 4, and 9 are briefly described below as they are not specific to massive RCT pathologies. Clusters 2, 5, 6, 7, and 8 are detailed later with their more RCT-associated pathogenic capabilities.

**Figure 2:**
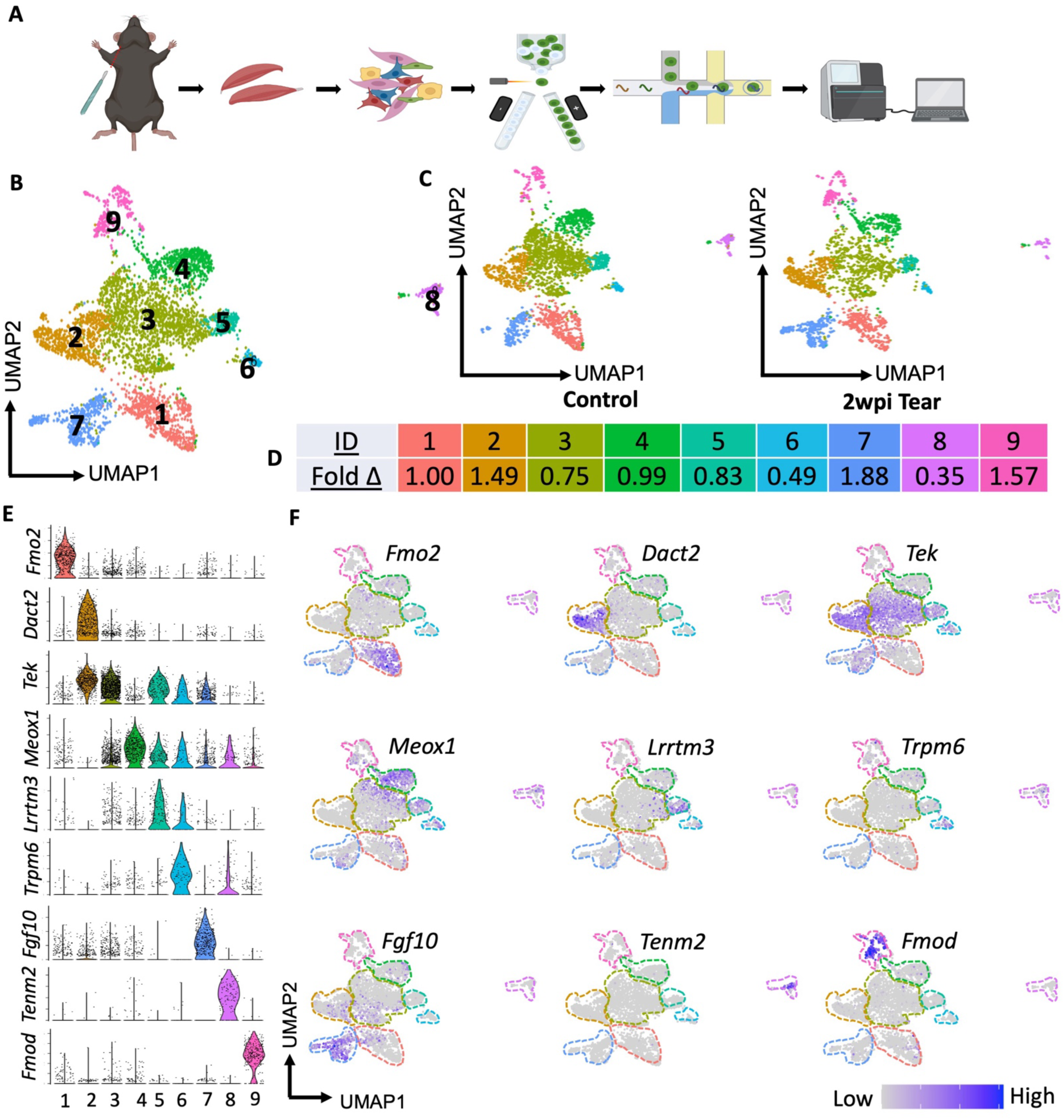
Control and 2wpi single cell RNA sequencing in murine rotator cuff. A) Schematic of scRNA data generation. B) Integrated (uninjured control and 2wpi tear) UMAP projection with 9 cluters. C) Separation of control and 2wpi UMAPs. D) Fold-change of cluster size between control and tear E) Violin plots of cluster specific gene markers. F) Corresponding feature plots of cluster specific gene markers. Schematics crafted with Biorender.

### Cluster 1 – *Osr1*+ Progenitor NMMCs

In rotator cuff muscles, the gene *Fmo2* exclusively marks Cluster 1 (***Fig. 2E,F***). This cluster is also a strong expressor of *Osr1,* which we and others have found defines the developmental and injury activated subpopulation of *Pdgfra+* NMMCs (37, 49–51). Prior work has indicated that these cells are progenitors to other NMMCs during early development (31, 32). Other work suggests *Osr1+* cells in the adult hindlimb are challenging to find because of extremely low transcript levels; however, arise quickly in response to injury (49, 50). Despite this we observe many *Osr1+* NMMCs in Cluster 1 of the uninjured and injured adult rotator cuff muscles (***Supp. Fig. 2A,B***). We confirmed their presence in adult mice using the *Osr1Cre^ERT2^;R26-Ai9^fx/fx^*mouse line. Many tdTOMATO+ cells are found interstitially within the SS muscles without injury (***Supp. Fig. 2D***). Following injury, the level of tdTOMATO+ cells increases (***Supp. Fig. 2D’***). Interestingly, this does not seem to be rotator cuff muscle specific since we also observe tdTOMATO+ cells in the uninjured tibialis anterior (TA) muscles, contrary to prior reports (50) (***Supp. Fig. 2E***). RNA velocity analysis of uninjured scRNA-seq data indicate that *Osr1+* cells move in a trifurcated trajectory toward Clusters 2, 5/6, and 7 (***Supp. Fig. 2C***).

### Cluster 3 – A Transitional Intermediate State or Population of NMMCs

Based on the depth of our scRNA-seq and trajectory analysis, Cluster 3 appears to be a transitional state or population between Cluster 2 and the other Clusters 4, 5, 6, and 8. There are no genes completely specific to this cluster, and most display a gradient of high gene expression in Clusters 2 or 4-6 that weakens with the transition across Cluster 3. A variety of genes marking these groups display a gradient of expression across Cluster 3. *Tek* (TIE1) is shown with high expression in Cluster 2 which spans all of Cluster 3, but reduces as the plot gets closer to Clusters 4-6, which have very low levels of *Tek+* cells (***Fig. 2 E,F***). Similar expression profiles are observed for *Dpp4, Cd55,* and several other genes (***Supp. Fig. 3A***). Genes such as *Meox1*, *Grm8,* and others all show opposing gradients with high expression in Clusters 4, 5, and/or 6 with an opposing gradient across cluster 3 (***Supp. Fig. 3B***). This NMMC subpopulation fate is hypothesized to be fluid and these cells likely represent those “transitioning” cells. This is further supported by our RNA velocity analysis showing trajectories in both directions across Cluster 3 (***Supp. Fig. 2C***). Following massive RCT, this subpopulation size is reduced, likely due to cells having gene expression profiles which resemble the other functionally relevant and/or more differentiated subpopulations (Cluster 2 or Clusters 4-6, and 8).

**Figure 3:**
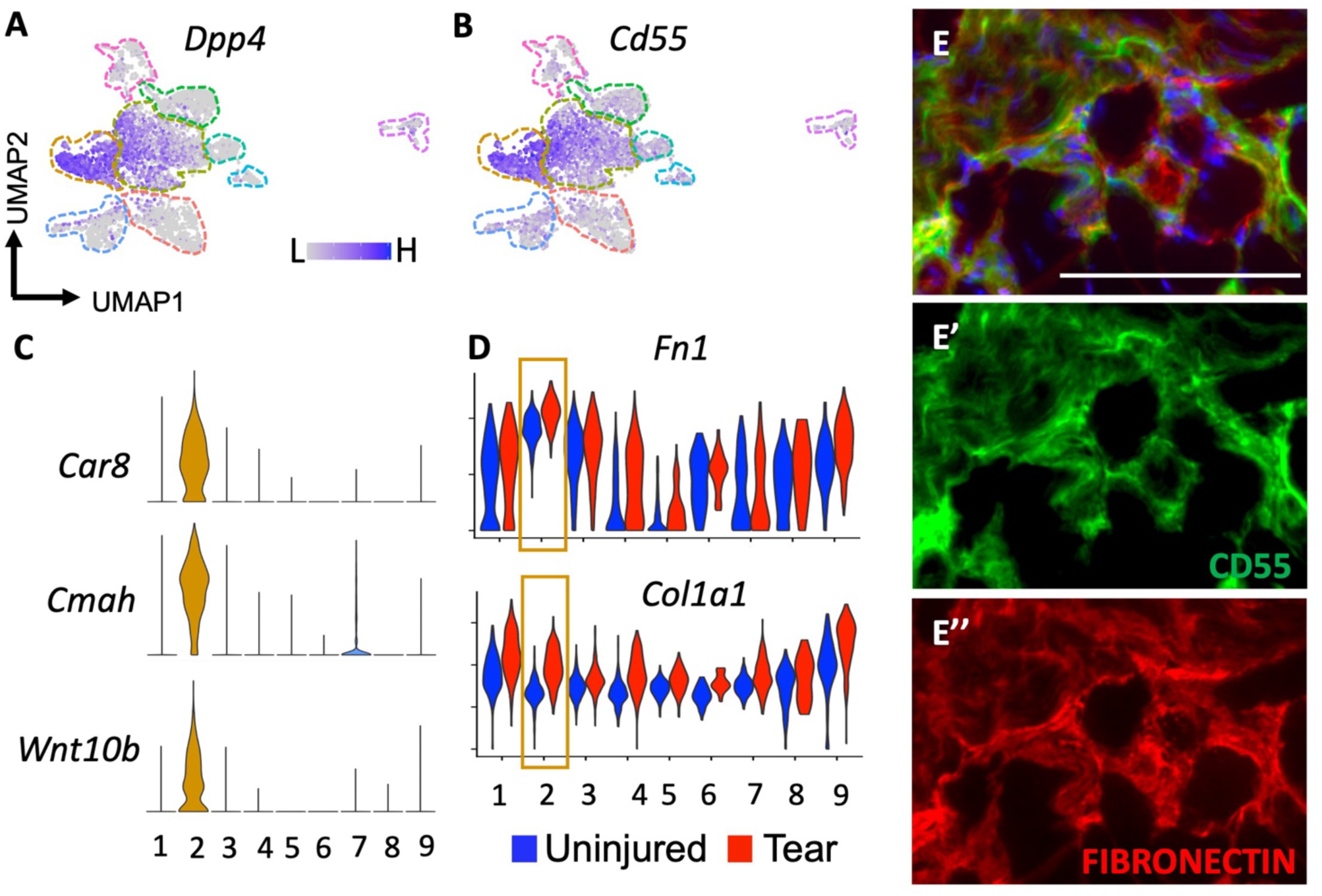
*Dpp4+*/CD55+ NMMCs contribute to massive RCT induced fibrosis. A) Feature plot of *Dpp4* on integrated UMAP. B) Feature plot of *Cd55* on integrated UMAP. C) Violin plots for Cluster 2 markers *Car8, Cmah,* and *Wnt10b.* D) Split violin plots showing expression increased in *Fn1* and *Col1a1* in cluster 2. E) 8wpi supraspinatus section showing colocalization of CD55 (E’) and FIBRONECTIN (E’’). All scale bars = 100um.

### Cluster 4 – *Meox1+* Mineralizing NMMCs

Cluster 4 contains cells which we have previously defined as mineralizing NMMCs (37). These NMMCs express genes such as *Clu* and *Hmcn1* (***Supp. Fig. 4C*)**, and are also more uniquely marked by *Ddit4l* and *Meox1* (***Fig. 2E,F, Supp. Fig. 4A,B*)**. Further, they express mineralization markers *Prg4* (52) and *Piezo2* (53). Cluster 4 cells do not have a population size or overt genetic response to massive RCT injury (***Fig. 2D***). This is likely because there are no documented mineralizing pathologies associated with RCTs in humans or mice (54).

**Figure 4:**
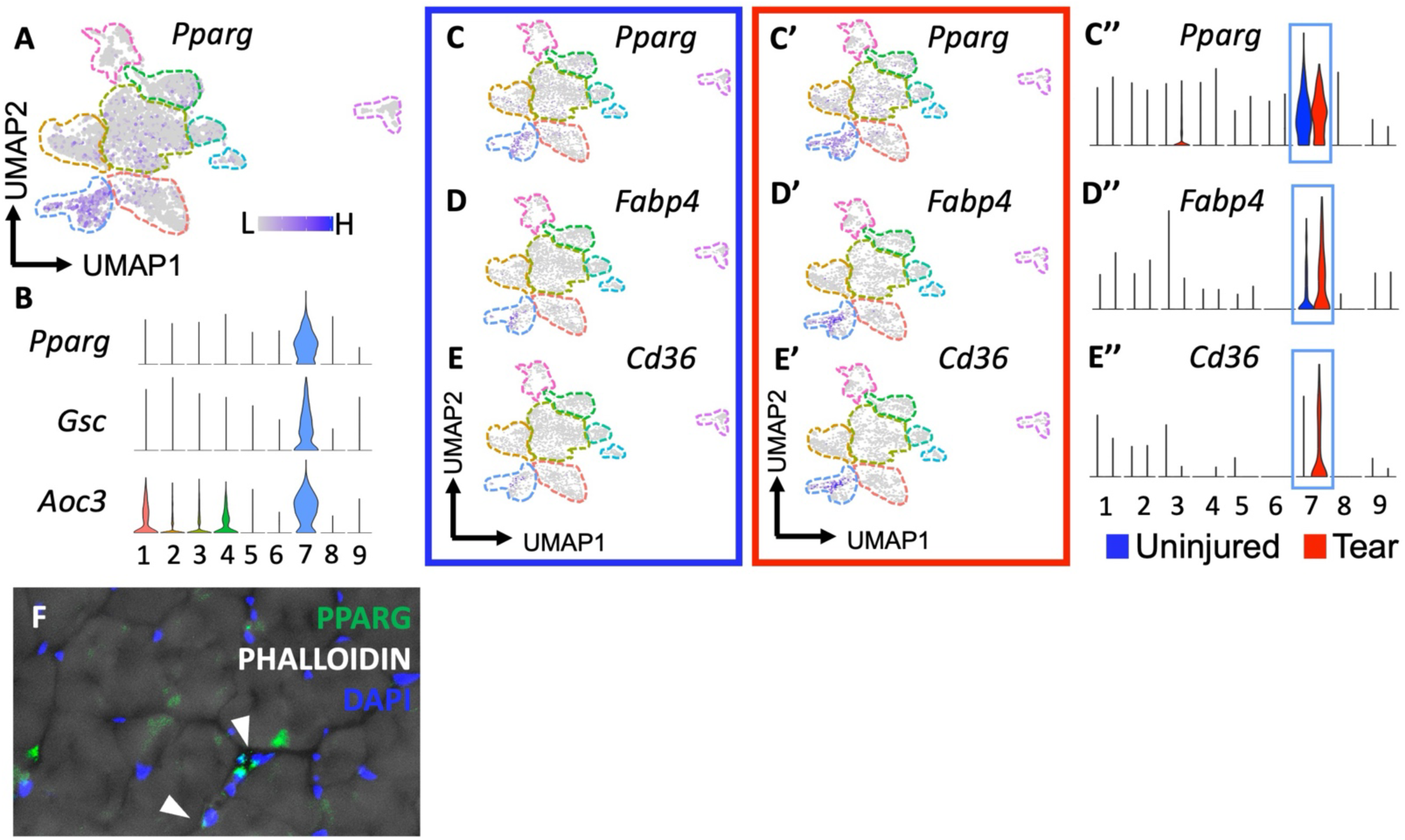
Homeostatic *Pparg+* NMMCs become more adipogenic after massive RCT. A) Feature plot of *Pparg* on integrated UMAP. B) Violin plots for Cluster 2 markers *Pparg, Gsc,* and *Aco3.* C) Increases in *Pparg* expression after tear with uninjured feature plot (C), 2wpi tear feature plot (C’) and split violin plot (C’’). D) Increases in *Fabp4* expression after tear with uninjured feature plot (D), 2wpi tear feature plot (D’) and split violin plot (D’’). E) Increases in *Fabp4* expression after tear with uninjured feature plot (E), 2wpi tear feature plot (E’) and split violin plot (E’’). F) Uninjured adult supraspinatus section showing PPARG+ NMMC nuclei interstitial in the PHALLOIDIN stained myofibers.

### Cluster 9 – *Tnmd*+ Tenocyte-like Tendon Progenitors

Cluster 9 is a group of *Fmod+* and *Tnmd+* tenocyte-like NMMCs (***Fig. 2E,F, Supp. Fig. 5A,C***). *Fmod* is a marker of tendon progenitors, and these TENOMODULIN+ cells are found interstitially in the body of skeletal muscles, distant from the myotendinous junction (37, 55, 56). Interestingly, they are usually classified as *Pdgfra-,* yet in the rotator cuff we find them to have lower transcripts; however, still maintain an identifiable amount of *Pdgfra* promoter induced GFP positivity and *Pdgfra* expression (***Supp. Fig. 5B***). In the rotator cuff, genes such as *Edil3, Col11a1,* and *Cilp2* are uniquely expressed in this cluster and play known roles in tendon, muscle, and skeletal extracellular matrices (ECMs) (55, 57) (***Supp. Fig. 5A***). Following massive RCT they respond by dramatically upregulating their *Tnmd* expression as well as expressing critical tenocyte-associated developmental transcription factors such as *Scx* and *Mkx* (***Supp. Fig. 5C*)**. These data suggest that Cluster 9 represents an endogenous population of tendon progenitors within skeletal muscle that are receptive to signals of RCT-induced tendon damage and may be important players in tethering the muscle to tendon ECM deep within the body of skeletal muscle.

**Figure 5:**
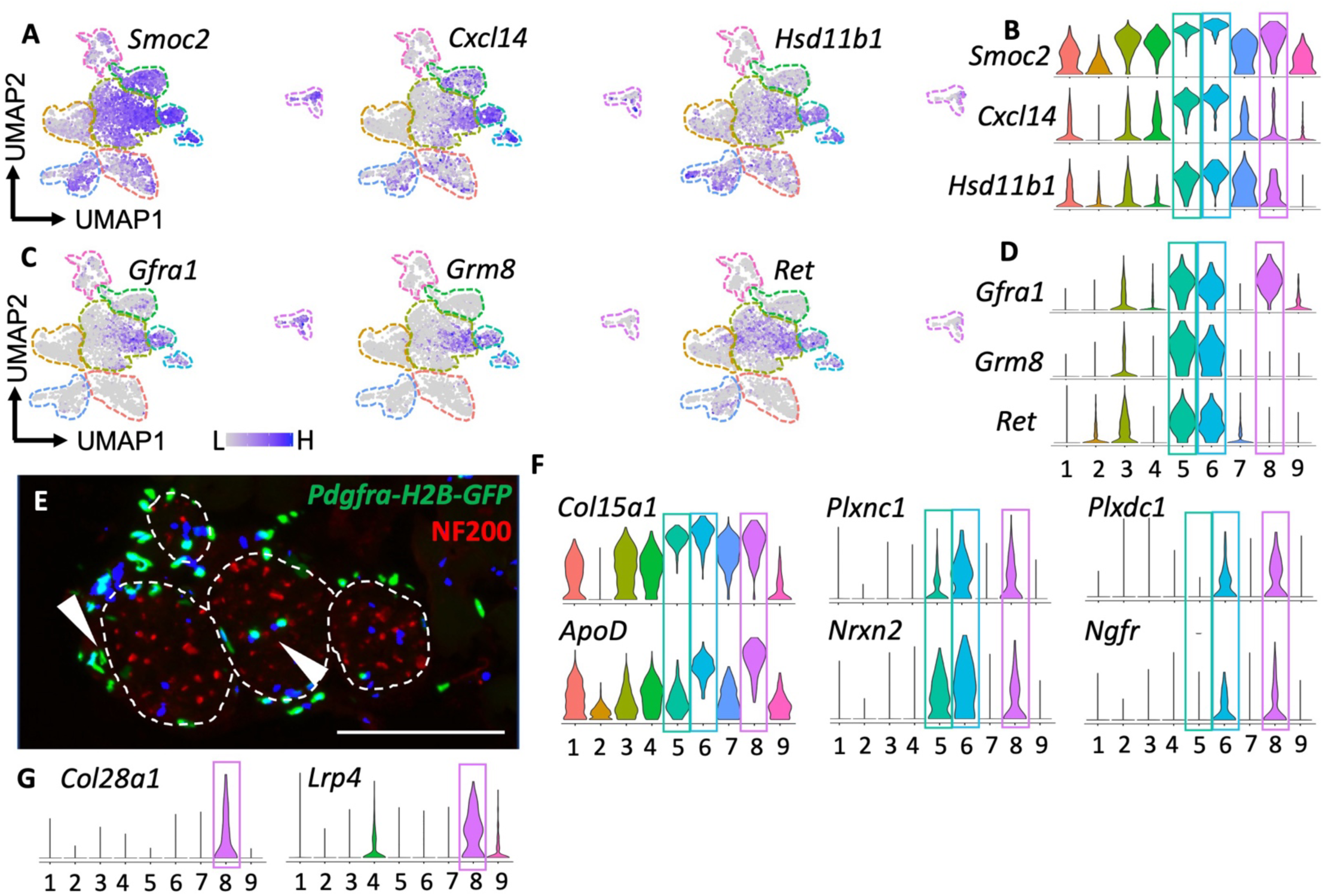
Clusters 5, 6, and 8 are Nerve Associated NMMCs. A) Feature plots of previously defined NMJ NMMC defined genes *Smoc2*, *Cxcl14*, and *Hsd11b1* on integrated UMAPs. B) Violin plots of *Smoc2*, *Cxcl14*, and *Hsd11b1*. C) Feature plots of additional nerve associated markers *Gfra1*, *Grm8*, and *Ret* on integrated UMAPs. B) Violin plots of *Gfra1*, *Grm8*, and *Ret*. E) Adult supraspinatus section from *Pdgfra-H2B-GFP* mouse showing intramuscular nerve cross sections (white outline, NF200 stained) with Pdgfra+/GFP+ nuclei sitting on the nerve sheath and within the nerve (white arrows). F) Violin plots of additional nerve/neuronal associated genes *Col15a1, Apod, Plxnc1, Nrxn2* expressed in these clusters. G) Violin plots of *Col28a1* and *Lrp4*. All scale bars = 100um.

### Cluster 2 – *Dpp4*+ Fibrogenic NMMCs

Cluster 2 represents a population of *Dact2+* and *Dpp4+* injury responsive NMMCs (***Fig. 2E,F, Fig. 3A***) (27, 37), which also express *Cd55, Pi16, and Tgfbr2* (***Fig. 3B, Supp. Fig. 6*)**. We identified *Wnt10b, Cmah,* and *Car8* to be uniquely expressed within this cluster (***Fig. 3C*)**. Following massive RCT, the Cluster 2 population of NMMCs increases 1.49-fold with an increase in the expression of fibrogenic markers, *Fn1* (58) and *Col1a1* (23, 59) (***Fig. 2D, Fig. 3D*)**. Based on these data, we hypothesized that the *Dpp4+/Cd55* subpopulation is responsible for RCT induced fibrosis and may become pathologic myofibroblasts secreting various ECM thickening/fibrosis inducing molecules. Using the 8wpi *PdgfraCre^ERT2^;R26-Ai9^fx/fx^* rotator cuff muscle samples, we identified that all FIBRONECTIN+ fibrotic plaques (which we’ve shown are TOMATO+) colocalized with CD55 immunostaining (***Fig. 3E*)**. These data indicate that the *Dpp4+/Cd55+* NMMCs (Cluster 2) are the likely source of the fibrotic ECM deposition following massive RCTs.

**Figure 6:**
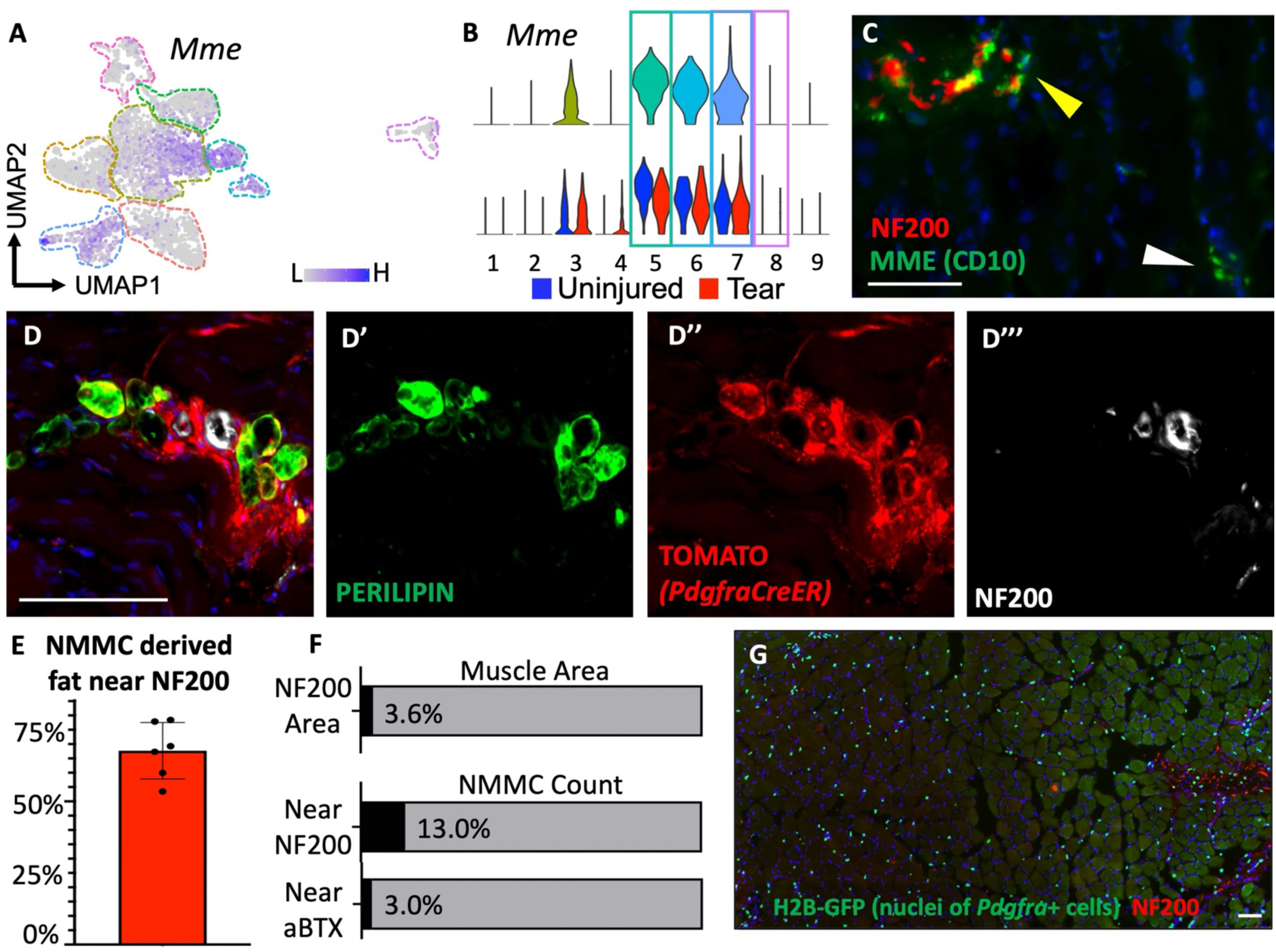
Nerve Associated NMMCs are responsible for majority of RCT induced fat. A) Feature plot of *Mme* on integrated UMAP. B) Integrated and split violin plot of *Mme.* C) Uninjured supraspinatus section showing MME+ staining near NF200+ (yellow arrow) and not associated with neural structures (white arrow). D) Representative image of PERILIPIN+/TOMATO+ fat near NF200 stain in 8wpi *PdgfraCreER;R26-Ai9^fx/fx^*supraspinatus sections. E) Quantification of localization PERILIPIN+/TOMATO+ fat near NF200 stain in 8wpi *PdgfraCreER;R26-Ai9^fx/fx^*supraspinatus sections. Near defined as touching or >3 myofiber cross sectional areas away from NF200+ stain. Data are represented as mean±SEM; n=6, N=3. F) Quantifications of NF200 area as a percent of total supraspinatus cross-section and then assessment of GFP+ nuclei near NF200 or aBTX (neuro-muscular junction) stain as a percent of all GFP+ cells in adult uninjured *Pdgfra-H2B-GFP*. Data represented as mean; n=4, N=3. G) Representative image of uninjured *Pdgfra-H2B-GFP* cross-section with small amount of NF200. All scale bars = 100um.

### Cluster 7 – Pre-adipogenic NMMCs

Cluster 7 represents a group of NMMCs that express the adipogenic regulators, *Fgf10* (60) and *Pparg* (61), under both normal and injury conditions (***Fig. 2E,F, Fig. 4 A,B***). These NMMCs also express the genes *Gsc* and *Aoc3* (***Fig. 4B***). *Pparg+* NMMCs are found in other muscles/developmental timepoints; however, are not typically found at a similar proportion as we have identified in the adult murine rotator cuff muscles. At 2wpi these NMMCs expand almost 2-fold (***Fig. 2D***) and upregulate their expression of the adipogenic master transcription factor, *Pparg,* as well as begin to express adipogenic differentiation genes, including *Fabp4* (62) and *Cd36* (63, 64) (***Fig. 4C-E***). Immunostaining of uninjured SS sections validates that PPARG+ nuclei are found interstitially between myofibers within the muscle (***Fig. 4F***). Based on these data, these NMMCs are a likely candidate for RCT-induced intramuscular fat deposition.

### Clusters 5, 6, and 8 – Nerve-associated NMMCs

Clusters 5, 6, and 8 represent nerve associated NMMCs. These subgroups of *Smoc2+, Hsd11b1+, Cxcl14+* cells respond to nerve damage (***Fig. 5A,B*)** and are associated with the capping of axons at the neuromuscular junction (30, 37); however, their specific localization/interaction with the myelinated nerve has not previously been described (though other Pdgfra+ cells can associate with nerves (65)). NMMCs in these clusters share expression of other neural genes such as *Gfra1, Nrxn2,* and *Grm8* (***Fig. 5C,D***). Additional nerve supportive and signaling genes such as *Apod, Nrxn2, Plxnc1, Ngfr,* and *Plxdc1* are localized to this cluster set (66–69) (***Fig. 5F***). Further, these subpopulations express collagens known to be in the peripheral nervous system (PNS), such as *Col15a1* which contributes to maturation of PNS nerves and C-fibers (70) (***Fig. 5F***). More specifically, Cluster 8 expresses *Col28a1* which surrounds non-myelinating glial cells and is found at nodes of Ranvier, where the myelin sheath is interrupted in myelinated fibers (71) (***Fig. 5G***). Cluster 8 also expresses genes related to NMJ localization and maintenance including *Lrp4* (72, 73) and *Tenm2* (74) (***Fig. 2E,F, Fig. 5G***). Uninjured adult supraspinatus muscle sections from the *Pdgfra-H2b-Gfp* mouse line indicate the presence of GFP+ nuclei along the edges of and within NF200 staining of myelinated nerves (***Fig. 5E, white arrows*)**. By 2wpi following massive RCTs, the overall cell numbers within these subpopulations decline dramatically (***Fig. 2D***).

To further confirm these cells are nerve associated we sought to integrate them with a public dataset of peripheral nerve fibroblasts (65). Nerve fibroblasts are *Pdgfra*-expressing cells in the PNS and play roles in PNS development, regeneration, and homeostasis, interacting with nerve axons and Schwann cells (75, 76). We downloaded these injured and uninjured data (GSE120678), subset for *Pdgfra*+ cells, and integrated with our own data subset on clusters 4, 5, 6, and 8. The resulting overlaid UMAP showed cells from our rotator cuff intermixed with the nerve fibroblasts in three larger groups (***Supp. Fig. 7***). These data provide strong evidence that clusters 4,5,6, and 8 from our RCT NMMCs are nerve associated cells.

**Figure 7:**
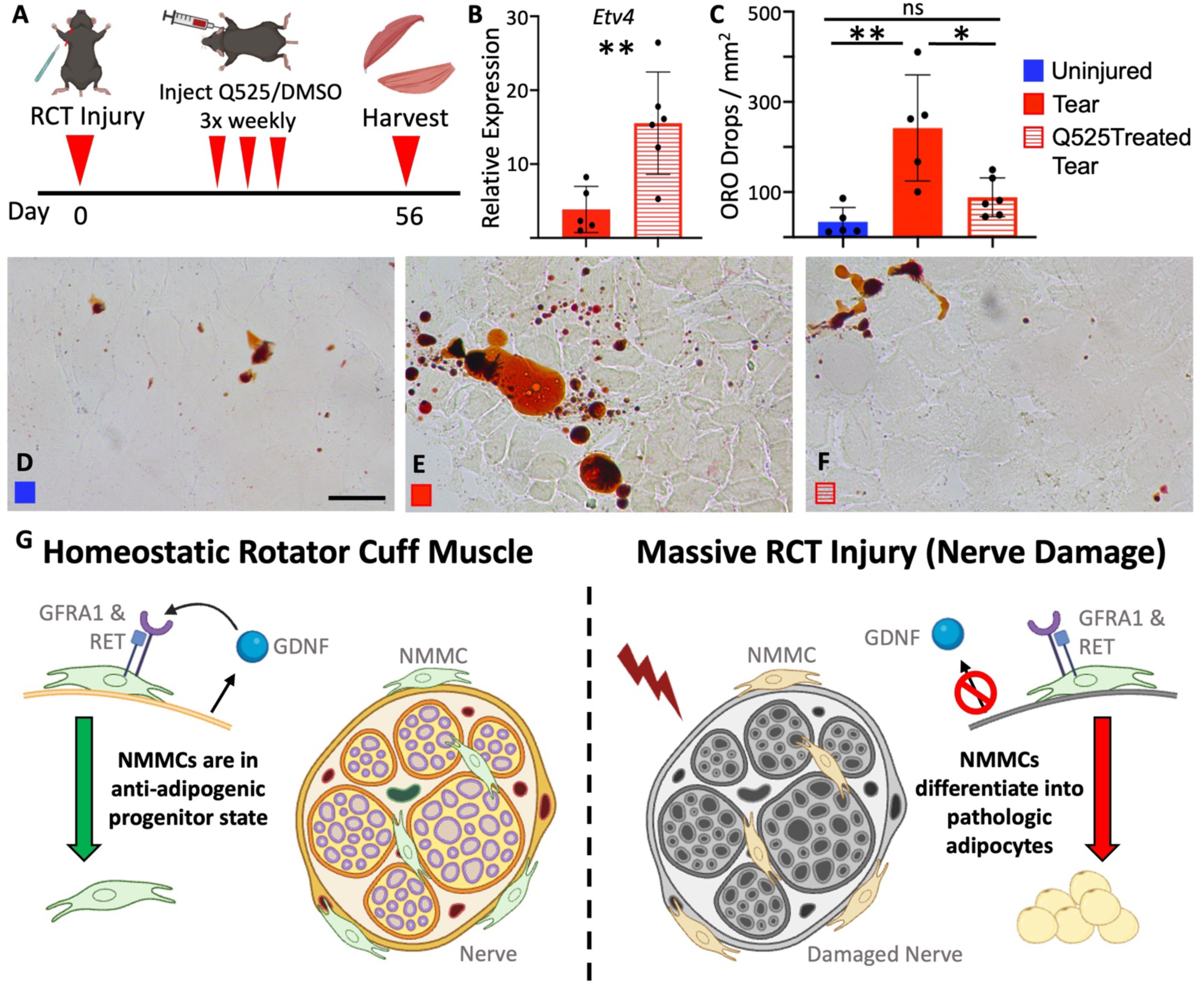
Treatment with RET agonist Q525 significantly reduces RCT induced fat. A) Schematic of injury and drug treatment. B) Whole muscle qPCR (infraspinatus) results of *Etv4* transcript to show downstream signaling from RET. Data are represented as mean±SD. n=5/6, N=2. Significance determined using unpaired student t-test, p=0.0072. C) Quantification of fat (oil red O) in supraspinatus sections from contralateral control, DMSO vehicle treated tear, and Q525 treated tear. Data are represented as mean±SEM. n=5/6, N=2. Significance determined using one-way ANOVA with multiple comparisons. Control vs. DMSO tear p=0.002, Control vs. Q525 tear p=0.439, DMSO treated tear vs. Q525 treated tear p=0.010. Representative images of oil red O staining for contralateral (D), DMSO treated tear (E), and Q525 treated tear (F). G) Proposed mechanism of GDNF-GFRA1-RET signaling axis keeping nerve associated NMMCs in progenitor state during homeostasis and how this is lost after nerve damage. Schematics crafted with Biorender.

### Nerve-associated NMMCs Preferentially Give Rise to RCT-induced Adipocytes

*Mme/*MME+ (CD10) NMMCs are found in several other muscle groups within both human muscle biopsies and murine muscles, while also being identified as a highly adipogenic subpopulation of NMMCs (39). Notably, expression of *Mme* is found not only in the Cluster 7 pre-adipocytes, but also within the nerve associated NMMC Clusters 5, 6, and 8 (***Fig. 6A,B***). Protein expression at both of these locations (interstitial to muscle fibers and near nerves) was confirmed *in vivo* (***Fig. 6C***). We sought to detangle the contribution of each of these subpopulations to the fatty infiltration pathology that is a hallmark of murine and human massive RCTs. To do so we assessed the localization of RCT-induced fat in the 8wpi *PdgfraCre^ERT2^; R26-Ai9^fx/fx^* lineage tracing muscle samples. Regeneration of the damaged SSN is complete by this time, and since nerve regeneration proceeds along the original nerve tracts (77), we determined the percent of fat which was localized near nerve structures. Using NF200 to mark the myelinated nerve structures, we stained our 8wpi RCT sections with NF200 and PERILIPIN and then assessed the number of PERILIPIN/TOMATO+ cells at or near (defined as 2 myofiber cross-sectional areas) the NF200+ area. We identified 67.7% (± 9.85) of the PERILIPIN/TOMATO+ adipocytes localize to nerve structures (***Fig. 6D,E***). Thus, nearly 70% of the RCT-induced fat resides near these nerve structures despite only 3.63% (± 4.85) of total muscle area being NF200+ and only 16.0% of *Pdgfra-H2b-Gfp* NMMCs residing near nerves (13.01%) or aBTX+ NMJs (3.01%) (***Fig. 6F***). Images of adult uninjured SS from *Pdgfra-H2b-Gfp* animals clearly detail the extent of *Pdgfra+* NMMCs localized interstitially to muscle fibers far from any identifiable nerve structures (***Fig. 6G***). Since most of the RCT-induced fat localized to nerve-associated regions in the mouse, these data indicate that this RCT-induced fat is likely generated from the nerve associated NMMCs and not the resident pre-adipocytes. It is of note that Clusters 6 and 8 are unique NMMC expressors of Hedgehog responsive genes, and blockade of Hedgehog signaling within NMMCs demonstrates reduced NMMC-associated adipogenesis in certain nerve/muscle injury settings (25, 78) (***Supp. Fig. 8***).

### GFRA1-RET-GDNF Signaling Axis is a Critical Regulator of RCT-induced Fatty Infiltration

Our scRNA-seq data indicate that many of the nerve associated NMMCs specifically express *Gfra1* and its co-receptor, *Ret* (***Fig. 5C,D***), which plays a role in regulating adipogenesis via GDNF signaling (79). Since motor neurons and their associated Schwann cells express GDNF (80), and the nerve associated NMMCs are in direct contact with or are within very close proximity to nerves and NMJs, we hypothesized that in uninjured contexts the SSN signals to the nerve associated NMMCs within the rotator cuff muscle through GDNF-GFRA1/RET, maintaining them in their normal homeostatic and supportive state. Upon RCT, which includes damage to the nerve, these nerve-associated NMMC signals cease, and the cells aberrantly differentiate into adipocytes. Therefore, we sought to maintain some level of GFRA1/RET signaling following nerve damage in our RCT model. To achieve this, we utilized a small molecule RET agonist, Q525 (81). The RET agonist or DMSO control was delivered via subcutaneous injection over the injured shoulder three times weekly from onset of injury until 8wpi (***Fig. 7A***). Injured and contralateral SS were harvested for histology and IS muscles were used for whole muscle qPCR. To validate Q525 activation of GDNF signaling in RCT muscles, we performed qPCR for *Etv4,* a downstream effector of GDNF-GFRA1/RET signaling. *Etv4* transcripts are over 5-fold higher in the Q525 treated RCTs compared to the DMSO treated RCTs (p=0.0072), indicating that the small molecule RET agonist, Q525, can penetrate the muscle and induce signaling (***Fig. 7B***). Next, we quantified fat levels in RCT and control muscles via Oil Red O staining. Here we show that Q525 treated RCTs have significantly lower levels of fat as compared to DMSO treated tears (***Fig. 7C-F***). Further, this 63% reduction in fat following Q525 treatment of RCTs was not significantly different than uninjured contralateral controls (***Fig. 7C-F*)**. Thus, we conclude that the significant increases in intramuscular fat induced via massive RCTs are due to nerve associated NMMC adipogenic differentiation and is at least partially mediated via the loss of GFRA1/RET signaling within these cells (***Fig. 7G***). Assessment of fibrosis using FIBRONECTIN immunostaining shows no changes in the levels of expression between Q525 treated and DMSO treated RCTs, with both exhibiting elevated pathologic levels as compared to uninjured controls (***Supp. Fig. 9***). These data are consistent with fact that GFRA1/RET receptors are not significantly expressed in the *Dpp4+/Cd55+* fibrogenic NMMCs (***Figs. 3A,B and 5C,D***) that are the primary contributors to RCT-induced fibrosis, and therefore the RET agonist treatment should have little to no direct impact on this population and RCT-induced fibrosis.

## DISCUSSION

Massive rotator cuff tears (RCTs) are one of the most common musculoskeletal injuries. Although some RCTs can be treated through physical therapy and surgical interventions, tendon retear rates are significant and have been correlated to pathologies such as intramuscular fat and fibrosis (1, 2, 4, 5, 11–13). No preventative measures or pharmacologic treatments exist for these pathologies as the cellular/molecular underpinnings have been previously unknown. In this study, we utilize a mouse model of massive RCTs to robustly detail the genetic heterogeneity of *Pdgfra+* NMMCs within the rotator cuff muscles and definitively demonstrate that these cells are the source of intramuscular fat and fibrotic ECM deposition and further define the functional roles of NMMC populations in massive RCT pathologies. Using lineage tracing, scRNA-sequencing, FACS, immunofluorescent staining, and RET agonist drug treatments, we identify a diverse heterogeneity of NMMCs that are also present in other skeletal muscles and demonstrate that specific nerve associated NMMCs are the primary cellular source of RCT-induced fat. Furthermore, this study opens new research avenues and identifies potential therapeutic targets to reduce RCT-induced pathologies to provide better surgical outcomes, while also promoting the exploration of nerve associated NMMC involvement in other adipogenic muscle disorders.

### Rotator Cuff NMMC subpopulations are like other muscle groups

Single-cell technologies in the last decade have uncovered genetic and proteomic heterogeneity in cell populations. These methods have been used in skeletal muscle biology in a few hindlimb muscles, at different developmental stages/ages and in the contexts of disease and injury. Despite this, our work is the first report of *Pdgfra+* NMMC heterogeneity in the rotator cuff muscles. This is important because RCT injuries develop a greater level of intramuscular fatty infiltration than most other tendon/muscle injuries. SS and IS muscles have a higher density of *Pdgfra+* NMMCs than other muscle groups. These cells are also more adipogenic *in vitro* and *in vivo* as compared to *Pdgfra+* NMMCs from other muscles, suggesting that their composition may be different (82). Finally, Pdgfra+ cells in the shoulder girdle come from both lateral plate mesoderm (like hindlimb) and neural crest (like craniofacial) lineages (83). While it is not assumed *Pdgfra*+ NMMCs from these distinct developmental lineages are functionally different, this aspect of their biology has never been dissected and thus remained a potential differentiator between rotator cuff *Pdgfra*+ NMMCs and NMMCs from other muscle groups before our work.

To achieve our goals, we performed deep sequencing of our single cell libraries (much deeper than other data sets; more than 150,000 reads/cell compared to ∼15,000-30,000 reads/cell from other studies), which allowed for more granular and detailed analyses that not only identified unique *Pdgfra*+ subpopulations, but also changes to important signaling molecules within these NMMC populations. These data defined subpopulations of *Pdgfra+* NMMCs found within the rotator cuff muscles that were similar and different than other muscle groups. Subpopulations included *Osr1+* developmental NMMCs, *Dpp4+* NMMCs, *Smoc2+* NNMCs, *Pparg+* NMMCs, and *Tnmd+* tenocyte-like NMMCs, which we further confirmed *in vivo.* Our study is the first report of OSR1+ cells in uninjured adult muscle and demonstrates robust rotator cuff *Osr1* promoter activity using the *Osr1Cre^ERT2^;R26-tdTomato* mouse line. Our inducible reporter data also show these cells are present in the uninjured adult TA muscle, which was previously thought to only turn on following injury. This disputed finding may stem from advances in the mouse genetic tools from the time the initial findings were published. We therefore determine that other muscles—not just the rotator cuff—possess a pool of the “progenitor/developmental” OSR1+/*Pdgfra+* NMMCs.

### Specific subpopulations play discrete roles in massive RCT pathologies

Through lineage tracing we determine that *Pdgfra+* NMMCs are responsible for the RCT-induced pathologies of intramuscular fatty infiltration and fibrosis. Fibrotic plaques specifically colocalize with CD55 cellular staining, implicating the Cluster 2 *Dpp4+/Cd55+* subpopulation of NMMCs in the massive RCT fibrosis pathology. This subpopulation has documented injury responsiveness following the regenerative muscle injury of BaCl_2_ injection into hindlimb muscles (27). Dpp4+ NMMCs are also necessary for the health of muscle resident macrophages (84)—further detailing their injury responsive qualities. Other published data indicate cytokine responsiveness to these cells and when Dpp4+ cells were harvested from skeletal muscle they were more adipogenic *in vitro* than other NMMC subpopulations, which is unlike their fibrotic RCT response in an atrophy setting such as RCTs. Interestingly, our data from the muscle atrophy RCT model shows population size increases at 2wpi; however, the same transcriptomic injury response in *Dpp4+* NMMCs is not seen this late in other regenerative injury models. This may be why in RCTs these NMMCs become pathologic myofibroblasts. Whether this is injury or muscle group specific remains unknown. Thus, Dpp4+ NMMCs are certainly injury responsive, but have differential responses based on injury and may function differently all together in *in vitro* experiments.

Regarding the adipogenic RCT-induced pathology, our data show nerve associated NMMCs are the primary cellular source of RCT-induced fat. The role nerve associated NMMCs specifically play in adipogenesis is previously unexplored. Fitzgerald. et al. determined MME+ cells in the hip muscle to be the most adipogenic subpopulation (39), but our scRNA-seq data provides a more robust characterization of *Mme* in the rotator cuff and shows expression in both pre-adipocytes and nerve associated NMMCs. Yet, in the context of massive RCT injury, the nerve associated NMMCs play a much greater role in adipogenesis and fatty infiltration. This may be the critical factor in understanding why rotator cuff tendon injury results in greater intramuscular fatty infiltration as compared to other tendon injuries (14). Though infrequently explored clinically, nerve involvement/damage has been indicated in rotator cuff injuries given the location of the SSN tract. After massive RCTs, SS and IS muscle retraction can induce damage to the nerve at both the suprascapular notch and the spinoglenoid fossa. Using histologic fat levels in human rotator cuff biopsies, Ibarra et al. confirmed a previous report (10) that full-tears have significantly more fat than control and high-grade partial tears, but further showed this significant increase is driven by increased fat levels from massive RCT patients—the group with the highest likelihood of nerve impingement. A small clinical report identified 8 of 8 patients with massive RCTs had EMG diagnosis of SSN neuropathy (85). Additionally, SSN nerve conduction velocity alterations have been observed in massive RCT patients and may be correlated with intramuscular fat development (46). Despite this, it is not current standard of care to perform EMG or NCV studies on RCT patients. In development of the murine model of massive RCTs, it was discovered that SSN damage was required to produce clinically relevant fat levels. Though initially a detractor of the model, our work highlights the importance of nerve involvement and its relevance to the human RCT pathology.

### Nerve associated NMMCs in other skeletal muscle injuries and diseases

Interestingly, RCTs may not be the only intramuscular adipose tissue (IMAT) forming injury for which nerve associated NMMCs contribute to the pathology. The classic skeletal muscle model of IMAT development, glycerol injections, are known to disrupt the NMJ. Specifically, this occurs at the pre-synapse, meaning it is damage to the end of the axon and not the receptors on the myofiber (86). In Kopinke et. al, the original paper describing *Pdgfra+* cells in skeletal muscle as the lineage of glycerol induced fat, the authors also genetically knocked out Hedgehog signaling components in *Pdgfra+* cells to reduce adipogenesis that was mediated via *Dhh* expression in endothelial and Schwann cells (25). Prior work also demonstrated that genetically removing cilia (hedgehog signaling inhibition) in NMMCs inhibited intramuscular adipogenesis following glycerol injury and in a mouse model of Duchenne muscular dystrophy (Dmd) (25). In our data, the Hedgehog signaling target genes are only found in nerve associated NMMCs (***Supp. Fig. 8***), lending to the conclusion they are the primary adipogenic contributing cells following glycerol injury and likely involved in the DMD pathology. Given the differential signaling mechanisms in glycerol-induced (regenerative-associated), DMD-induced (regenerative-associated), and RCT-induced (atrophy-associated) adipogenesis, it is likely that injury/disease context also plays an important functional role in controlling outcomes. Yet, it is possible that nerve-associated NMMCs play a significant role in multiple kinds of IMAT development. Therefore, in the future it will be important to explore this subpopulation in the contexts of other injuries/diseases that have fatty pathologies, like the variety of muscular dystrophies, ALS, and disuse.

Our comparative scRNA-seq analyses between RCT NMMC clusters and peripheral nerve fibroblasts further demonstrated that cluster 4 (*Meox1+/Clu+)* from our dataset is a nerve associated NMMC population. These *Clu*+ NMMCs have been previously defined as highly mineralizable NMMCs responsive to BMP signaling (37), which could be linked to heterotopic ossification (HO)—a disease of soft tissues such as muscle, tendons, and ligaments generating ectopic bone. HO has repeatedly been associated with neurotraumas including traumatic brain injuries, spinal cord damage, and sciatic nerve damage in routine hip arthroplasty (87–89). Additionally, overexpression of *Bmp4* in PNS neurons (not normal homeostatic Bmp4 producers) can induce HO, establishing a potential connection between mineralizing NMMCs (which can respond to BMPs) and nerve association (90). These data reflect the broad and important implications of our findings to not only RCT pathologies, but also muscle pathologies associated with HO, ALS, and other muscle diseases.

A growing body of work continues to uncover a genetic and functional heterogeneity of NMMCs contributing in specific and meaningful ways to skeletal muscle biology, disease, and injury repair/regeneration. Here we detail this heterogeneity for the first time in rotator cuff muscles in the presence and absence of massive RCTs. Due to the scRNA-seq depth of our study we were not only able to define this NMMC heterogeneity but were also able to uniquely identify signaling pathways and potential druggable targets responsible for the massive RCT pathologies. We determined that nerve associated NMMCs contribute to the majority of RCT-induced fat and is moderated at least partially via a GDNF-GFRA1-RET signaling axis. Currently, no clinical preventative measures or treatments for the RCT-induced pathologies exist, thus our work is a critical step in developing therapeutics for this common musculoskeletal injury. This work may also provide insights into other tendon injuries and musculoskeletal biology in general, especially for injuries/disease which result in intramuscular fat. Finally, it demonstrates a novel approach to using scRNA-seq technologies to further define pathogenic cell populations in order to gain better insights into molecular signaling dynamics with the goal of accelerating the identification and testing of therapeutic targets.

## METHODS

### Sex as a biological variable

This study examined both male and female animals with similar findings reported for both sexes. Lineage tracing and single cell RNA sequencing were performed on sex matched cohorts. Q525 drug treatment used male mice only.

### Mouse Lines

The *PdgfraCre^ERT2^* mice (JAX Strain 03277)*, R26-Ai9* or *R26-tdTomato* (JAX Strain 007909), *Pdgfra-H2b-Gfp* (JAX Strain 007669)*, Osr1Cre^ERT2^* (JAX Strain 009061), and wild-type *C57BL/6J* (JAX Strain 000664) mice were obtained from Jackson Laboratories. Both sexes of mice were used in this study, and they were housed at 23°C on a 12-h light/dark cycle and maintained on a PicoLab Rodent Diet. All animal work was approved by the Duke University Institutional Animal Care and Use Committees (IACUC). No sex differences were noted. All experiments except RET agonist (Q525) treatment experiments were performed on both male and female animals in sex matched cohorts; Q525 treatment was performed with male mice only.

### Massive rotator cuff tear injury (TT+SSN)

Animals were put under anesthesia using isoflurane. The upper left quadrant of the chest and left shoulder were shaved then washed 3 times alternating with 70% EtOH and Iodine solution. Animals were moved to a warming pad to maintain body temperature during surgery and draped. All surgical injuries were performed under a stereoscope. To perform the injury a 1cm incision was made over the clavicle and head of the humerus. The deltoid muscle was split to expose the humoral head, after which the supra- and infraspinatus tendons were transected (TT). After TT the SSN was exposed by moving the clavicle down and transected at the point the nerve descends from the brachial plexus. Following surgery, topical anesthetic and antibiotics were added to the closed incision and a wound clip was used to close the site. Animals recovered from anesthesia under observation. Injured animals were housed together (sexes kept separate) and monitored daily for 7 days post injury. After 10 days wound clips were removed.

### Tamoxifen injection (Lineage tracing assessments)

3-month-old mice were IP injected with tamoxifen for 5 days at a concentration of 100mg tamoxifen per 1kg of mouse weight using a 25-gauge needle. Mice were allowed to recover for at least 1 week before massive RCT injury.

### Q525 (Ret Agonist) injection

The small molecule Ret agonist (Q525) was diluted in DMSO and DMSO alone was used for control injections. Mice were anesthetized using isoflurane and 25ug of Q525 or DMSO was injected subcutaneously using a 25-gauge needle over the injured shoulder muscles. Care was taken to ensure the needle did not penetrate the muscle to ensure no regenerative injury occurred. Injections were performed 3-times per week starting at the time of injury with the last injection occurring 4 hours prior to 8wpi tissue harvest.

### Tissue harvest and histology preparation

Freshly isolated SS and IS muscles were rinsed in 1X PBS and incubated overnight at 4°C in 30% sucrose. Muscles were embedded in OCT matrix and flash frozen in isopentane, supercooled in a liquid nitrogen bath. Cryosections were cut at 15uM or thick 50uM sections. Cross-sectional tissue sectioning began at the end of the myotendinous junction and went throughout the length of the muscle.

### Immunostaining

Slides were brought to room temperature and dehydrated in PBS. Slides were post-fixed in 4% PFA and antigen retrieval (if necessary) was performed. Slides were blocked in 3% bovine serum albumin for 1 hour, then incubated in primary antibody overnight at 4°C in a humidified chamber. Slides were washed in PBS and incubated with secondary antibody for 45 minutes at room temperature. All antigen retrieval, primary antibody, and secondary antibody details are found in Table 1. Sections were mounted with ProLongGold Mountant with DAPI and imaged on a Leica DMI3000B microscope using a Leica DFC3000G camera with the latest Leica imaging suite or Zeiss Axio Imager.Z2 with the latest Zen imaging suite. Images were reconstructed and pseudocolored on Fiji ImageJ.

### Whole muscle qPCR

Harvested muscle was flash frozen in liquid nitrogen. Whole muscles were pulverized using a Retsch mixer mill MM400 and pulverized tissue was removed from the shakers using Trizol. RNA was extracted with bromo-chloro-phenol using phase lock tubes and a Quigen RNA isolation mini-kit. RNA concentration and purity were determined via Nanodrop metrics and cDNA prepped using an iScript kit. qPCR reactions for select genes were run in duplicate or triplicate for all samples on a BioRad CFX96 Real-Time System or ABI QuantStudio 3 Real-time PCR System.

### Oil Red O Staining and Quantification

Slides were brought to room temperature and dehydrated in PBS. Slides were post-fixed in 4% PFA then washed in PBS. Following PBS washes, the slides were dipped in water, 60% isopropanol, and incubated in oil red O (ORO) solution for 15 minutes. Following ORO they were again dipped in 60% isopropanol, water, then mounted with aqueous mounting media. Sections were imaged on a Leica DM2000 LED microscope using a Leica DFC450 camera with the latest Leica imaging suite. Images were stitched, ORO droplets quantified, and total muscle cross-sectional area determined using Fiji ImageJ software.

### H&E Staining

Slides were brought to room temperature and dehydrated in PBS. Slides were post-fixed in 4% PFA then washed in PBS. They were then stained with alcoholic eosin and counterstained with Mayer’s hemotoxylin. Slides were dehydrated through changes of ethanol and xylene, then mounted with cytoseal and imaged on a Leica DM2000 LED microscope using a Leica DFC450 camera with the latest Leica imaging suite.

### Picrosirius Red Staining

Slides were brought to room temperature and dehydrated in PBS. Incubated in Bouin’s fixative for 1hour at 60°C, washed with diH_2_O, and then incubated at room temperature in picrosirius red solution for 1 hour. Slides were then washed with Solution C, dehydrated in the ethanol/xylene gradient, and mounted with cytoseal. Slides were imaged on a Leica DM2000 LED microscope using a Leica DFC450 camera with the latest Leica imaging suite.

### Muscle resident cell isolations

Muscle interstitial cells from supraspinatus, infraspinatus, and various hindlimb muscles were released as described. Muscles were isolated and mechanically minced, then enzymatically digested in PBS with 10% FBS, 2mg/mL type II Collagenase, and 4mg/mL Dispase for 30-60 minutes rotating at 37°C. Sample tubes were shaken every 10 minutes during the digestion. Following digestions, samples were put on ice and the enzymatic reaction quenched with 2.5mL FBS.

### Flow Cytometry

Single muscle interstitial cells were isolated as described above for tear and contralateral control muscles (kept separate) from 5dpi, 2wpi, and 4wpi *Pdgfra-H2b-Gfp* mouse rotator cuff muscles. Red blood cells were lysed with ACK lysis buffer, then stained with DAPI for 30 minutes on ice. Cells were washed with PBS+10%FBS, spun down, and resuspended in PBS+2%FBS for flow cytometry analysis. Using a FACSCanto Analyzer cells were gated on GFP and DAPI to yield single live GFP+ cells. Percentage of live GFP+ cells to total cells were calculated using FlowJo V9 software to remove non-cellular events (debris), doublets, and DAPI+ cells.

### Single-cell isolation and scRNA-sequencing

#### Uninjured: rotator cuffs from 4 male and 4 female mice; Tear: 2wpi rotator cuffs from 2 male and 2 female mice

Single muscle interstitial cells were isolated as described above for tear muscles and uninjured control muscles (kept separate). Red blood cells were lysed with ACK lysis buffer, then stained with DAPI for 30 minutes on ice. Cells were washed with PBS+10%FBS, spun down, and resuspended in PBS+2%FBS for fluorescent activated cell sorting (FACS). Cells were gated on forward side scatter, side scatter, GFP, and DAPI, to yield single live GFP+ cells. Cells were taken to the Duke Molecular Physiology Institute’s Molecular Genomics core where single cells were isolated from highly viable cells, washed and filtered through a 40µm Flowmi Cell Strainers and resuspended in a 1x PBS / 0.04% BSA (+RNase inhibitor for nuclei) buffer.

A Cellometer Automated Cell Counter using AOPI fluorescent dye was used to determine the concentration of the single cell suspension and the viability of cells. Cells were then combined with a master mix that contained reverse transcription (RT) reagents. This master mix was loaded onto the 10x microfluidics chip, together with gel beads carrying the Illumina TruSeq Read 1 sequencing primer, a 16bp 10x barcode, a 12bp unique molecular identifier (UMI) and a poly-dT primer, and oil for the emulsion reaction. Using a 3’ next GEM v3.1 kit, the 10x Genomics Chromium X instrument used the microfluidics in the chip to partition the nuclei into nanoliter-scale gel beads in emulsion (GEMs) within which the RT reaction occured; all cDNAs within a GEM, created from a single cell, share a common barcode. After the RT reaction, the GEMs were broken, full length cDNAs cleaned with Silane Dynabeads, and then amplified via polymerase chain reaction (PCR) and purified using SPRIselect bead size selection. Resulting cDNAs were assayed using a 4200 TapeStation System High Sensitivity D5000 ScreenTape and reagents for qualitative and quantitative analysis.

Enzymatic fragmentation and size selection were used to optimize the cDNA amplicon size for the sequencing library preparation in which Illumina P5 and P7 sequences, a sample index, and TruSeq read 2 primer sequence are added through End Repair, A-tailing, Adaptor Ligation PCR. The final libraries contained P5 and P7 primers used in Illumina bridge amplification. Following additional bead purification, libraries were assayed for quality using 4200 TapeStation System HSD1000 ScreenTape and reagents, then quantitated and checked for successful adapter ligation with the KAPA Library Quantification Kit on the Applied Biosystems QuantStudio. Sequencing was performed by the Duke Genomic and Computation Biology Core using paired end sequencing on an Illumina NovaSeq6000 SP flow cell with 28×10×10×90 reads.

##### Computational analysis of scRNA-seq data

###### Unsupervised clustering and UMAP generation (rotator cuff)

The primary analytical pipeline for the scRNA-seq analysis followed the recommended protocols from 10X Genomics. Briefly, we demultiplexed raw base call (BCL) files generated by Illumina sequencers into FASTQ files, upon which alignment to the mouse reference transcriptome (with an artificial chromosome containing the *H2b-eGfp* sequence), filtering, barcode counting, and UMI counting were performed using the most current version of 10X’s Cell Ranger software. Using the Seurat platform in R each dataset (uninjured and tear) underwent individual basic filtering for minimum gene and cell observance frequency cut-offs (200-8000 genes for uninjured, 200-7500 genes for tear; 7% mitochondrial genes, genes expressed in >10 cell for both). The datasets were normalized and then integrated using Seurat’s FindAnchors() function. We then closely examined the data and performed further filtering based on a range of metrics in attempt to identify and exclude possible multiplets. We also thresholded *eGFP* on express of 0.2 or high to exclude the few cells which were FACS sorted, but were *eGFP* negative by gene expression and not FAPs by other marker gene expression.

After quality control was complete, we performed linear dimensional reduction, calculating principal components using the most variably expressed genes in our dataset (3000 variable genes, dims = 40). Significant principal components for downstream analyses were determined using the most variably expressed genes in our dataset. Cells were groups into an optimal number of clusters for de novo cell type discovery using Seurat’s FindNeighbors() and FindClusters() functions (resolution = 0.4), graph-based clustering approach with visualization of cells being achieved through the use of UMAP, which reduced the information captured in the selected significant principal components to two dimensions. Differential expression of relevant cell marker genes was visualized on UMAP feature and violin plots to reveal specific cell subtypes and tear induced changes.

###### Unsupervised clustering and UMAP generation (integration with GSE120678 data)

Cells from rotator cuff clusters 4, 5, 6, and 8 were subset from the data. Uninjured and 9dpi axotomy datasets were downloaded from GEO and subset on GFP+ cells. QC was performed as stated above and all data was integrated, grouped into optimal number of clusters for de novo cell type discovery using Seurat’s FindNeighbors() and FindClusters() functions (resolution = 0.3), graph-based clustering approach with visualization of cells being achieved through the use of UMAP, which reduced the information captured in the selected significant principal components to two dimensions.

###### RNA Velocity trajectory analysis

RNA velocity was computed using velocyto.py and velocyto.R (91). The aligned BAM file was processed using velocyto.py to obtain counts of unspliced and spliced read in loom format. The loom file was then processed using velocyto.R ub combination with an R package SeuratWrappers. Twenty nearest neighbors in slope calculation smoothing were used for RunVelocity.

### Statistics

The data were analyzed using Graphpad Prism 9 software. For comparisons between two groups 2-tailed t tests were used. For 3 or more groups, 1-way ANOVA with Tukey’s multiple-comparison tests were used. For histologic/immunofluorescent quantification data 2-3 replicates were averaged for each animal value. qPCR data were 2-3 technical replicates. A *p-*value of less than 0.05 was considered significant.

### Study Approval

All animal work was approved by the Duke University Institutional Animal Care and Use Committees (IACUC) under protocol number A037-23-02.

### Data Availability

All quantifications for figures and conclusions in the paper are present in supplemental materials. The scRNA-seq data is available on GEO via Accession ID GSE308502. Code will be made available upon acceptance of paper.

## AUTHOR CONTRIBUTIONS

HR: assisted in experimental design, conducted experiments, acquired data, analyzed data, and wrote and edited the manuscript; AJM, APL, JI: conducted experiments, acquired data, analyzed data; JVC: assisted in experimental design, analyzed data, and edited manuscript; MJH: designed research studies, analyzed data, wrote and edited the manuscript.

## ACKNOWLEDGEMENTS

We would like to acknowledge the assistance of the Duke Molecular Physiology Institute Molecular Genomics core for the generation of single cell RNA-seq libraries and the Duke Center for Genomic and Computational Biology for the sequencing of the scRNA-seq libraries. We would also like to acknowledge the Duke Light Microscopy Core Facility for their support with imaging and analysis, as well as, the Flow Cytometry Shared Resource Core at Duke for FACS and access to flow cytometric analysis. We would also like to acknowledge Dr. Jianhong Ou from the Duke Regeneration Center (DRC) for bioinformatics support with scRNA-seq data analysis. This research was supported in part by the following United States National Institutes of Health grants: R21 grant (AR083041 to MJH), F31 grants (AR076180 to APL and AR082701 to HR), T32 grant (HD040372 for support of HR), Precision Genomics Collaboratory-OBGE Graduate Student Pilot Research Grant Award from Duke University to HR, and departmental funds to MJH from the Department of Orthopaedic Surgery at the Duke University School of Medicine.

---

**Supplemental Figure 1:**
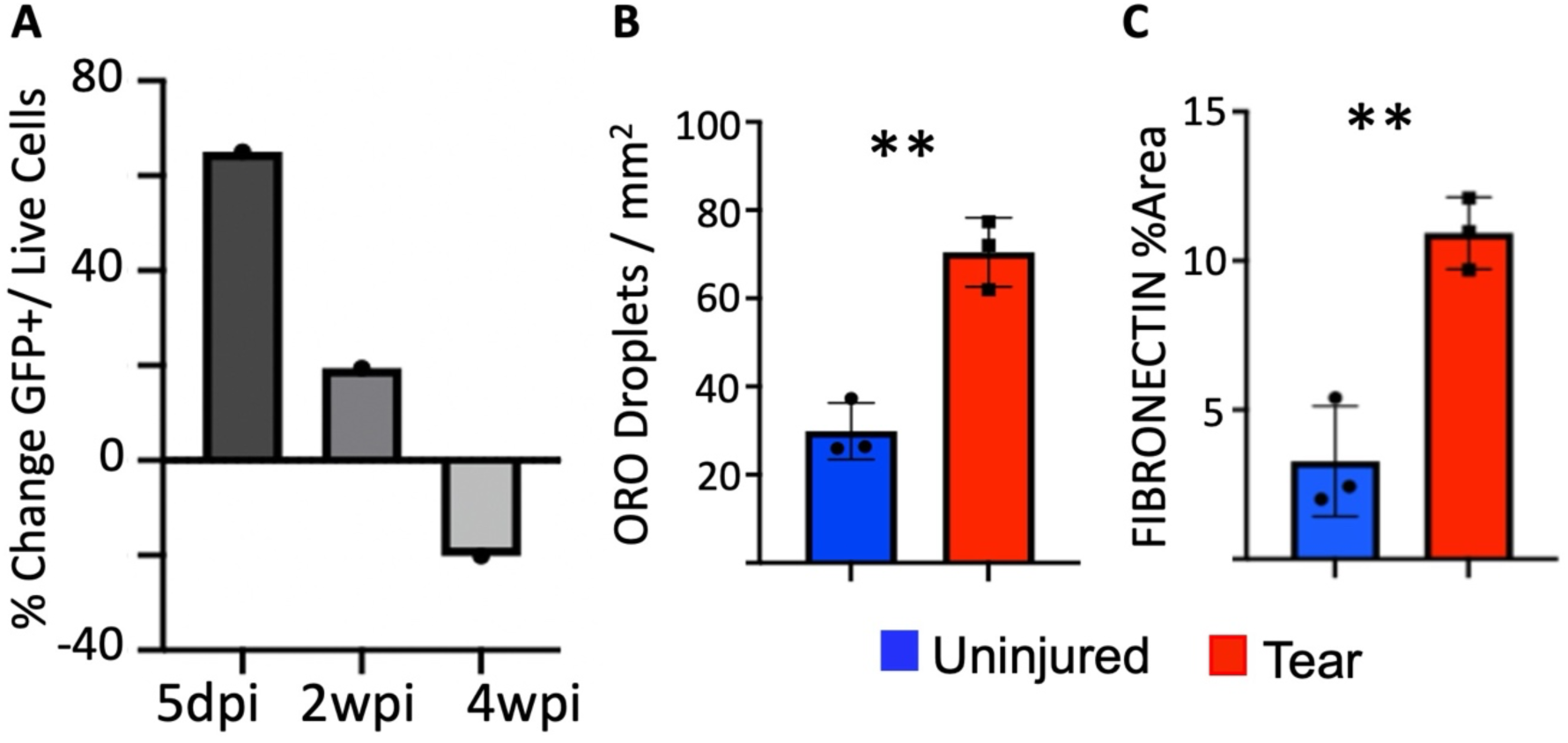
Justification of 2wpi for single cell. A) Percent change of GFP+ cells from tear to contralateral control in 5dpi, 2wpi, and 4wpi *Pdgfra-H2B-GFP* from digested SS and IS muscles. Data represented as means. B) Oil Red O staining quantification from 2wpi SS sections. Data are represented as mean±SD, p=0.006; n=3. C) FIBRONECTIN % Area quantification from 2wpi SS sections. Data are represented as mean±SD, p=0.008; n=3.

**Supplemental Figure 2:**
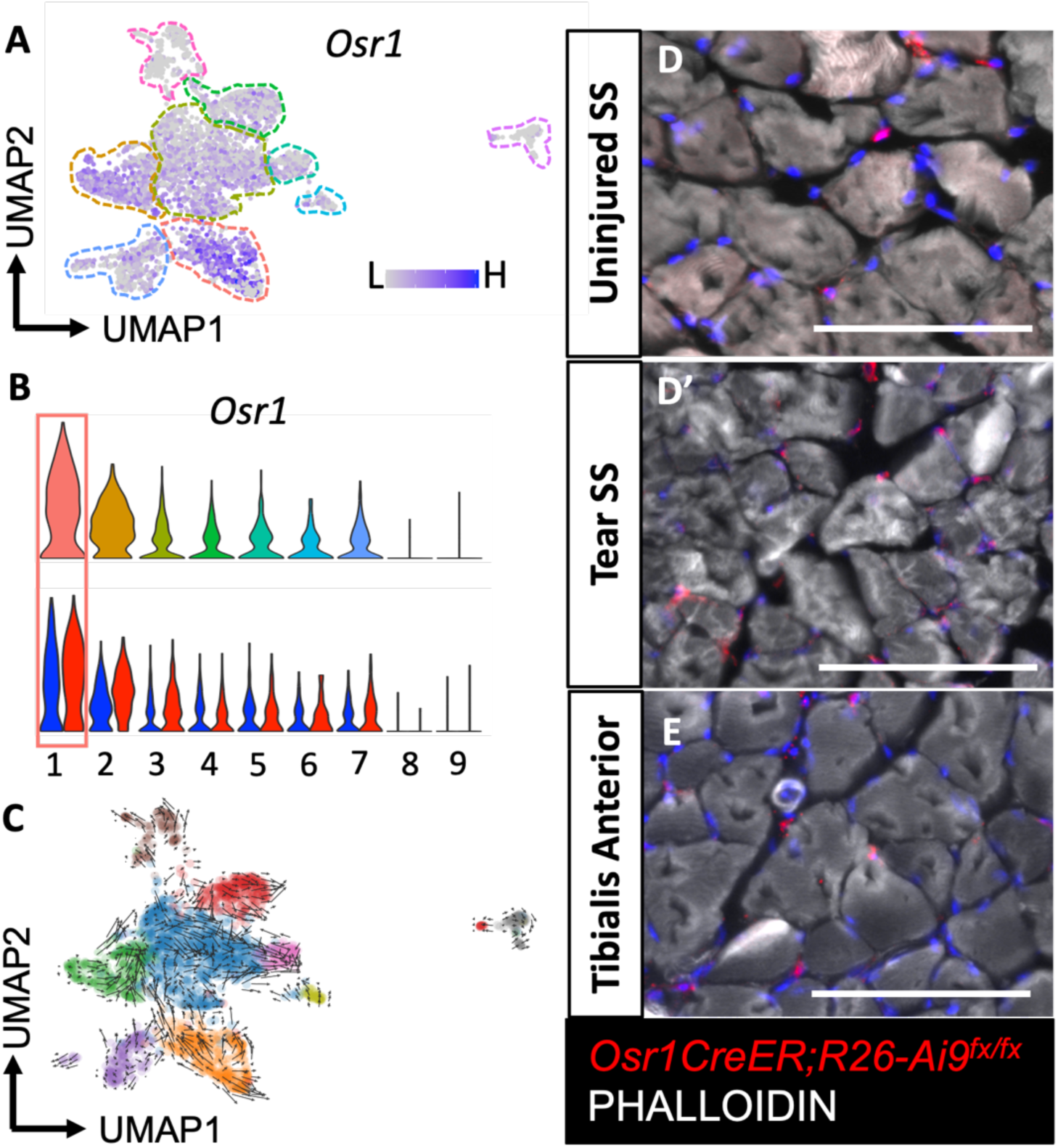
*Osr1+* progenitor NMMCs are found in adult rotator cuff. A) Feature plot on integrated UMAP, (B) violin plot, and split violin plot for *Osr1.* C) RNA Velocity of uninjured cells showing trajectory from Osr1+ cells. TOMATO+ cells from adult *Osr1CreER;R26-Ai9^fx/fx^* mice in uninured (D) and tear (D’) SS sections. (E) Adult tibialis anterior muscles in *Osr1CreER;R26-Ai9^fx/fx^* mice.

**Supplemental Figure 3:**
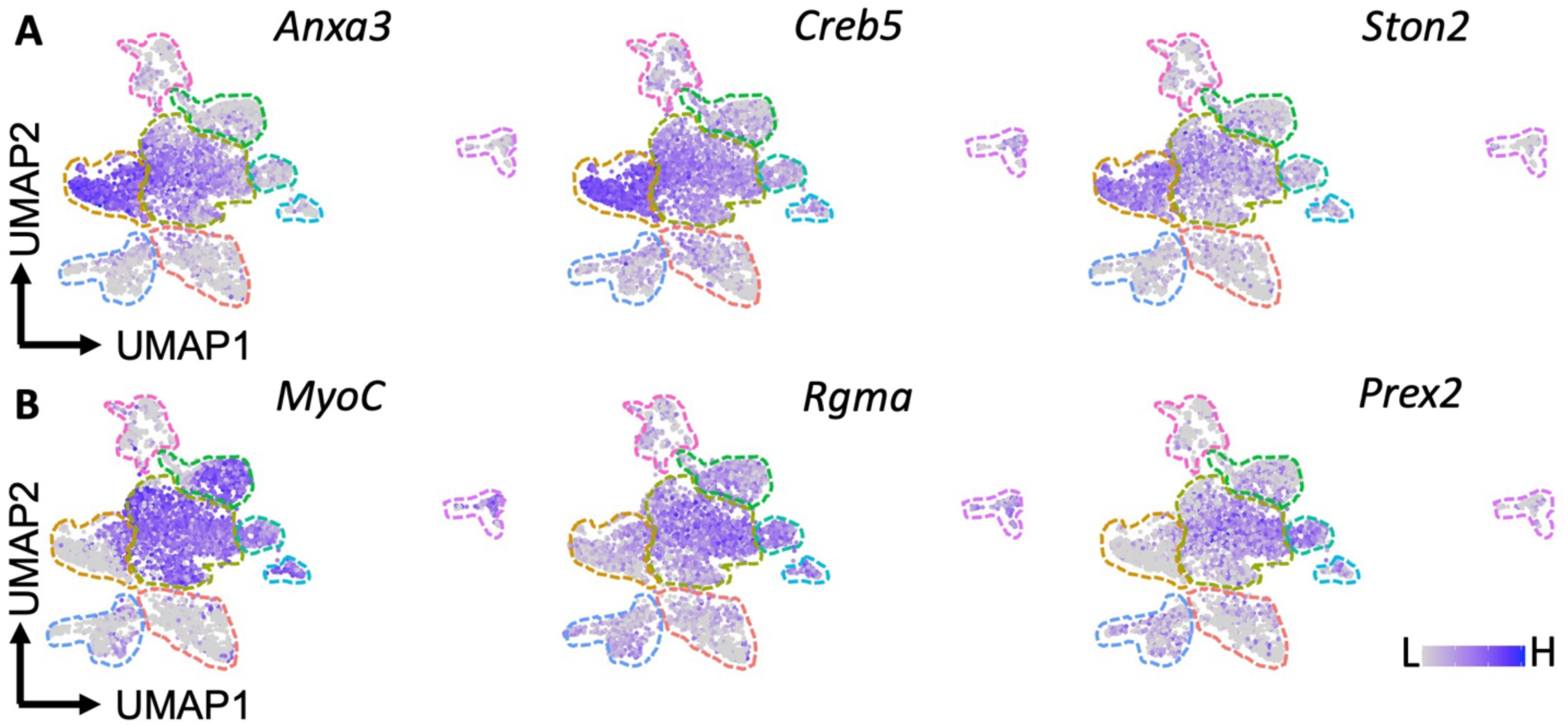
Cluster 3 is a transition population between Cluster 2 and Clusters 4, 5, 6, and 8. (A) Feature plots of *Anxa3, Creb5,* and *Ston2* which gradient high in Cluster 2 across Cluster 3. (B) Feature plots of *MyoC, Rgma,* and *Prex2* which gradient high in Clusters 4, 5, 6, and 8 across Cluster 3

**Supplemental Figure 4:**
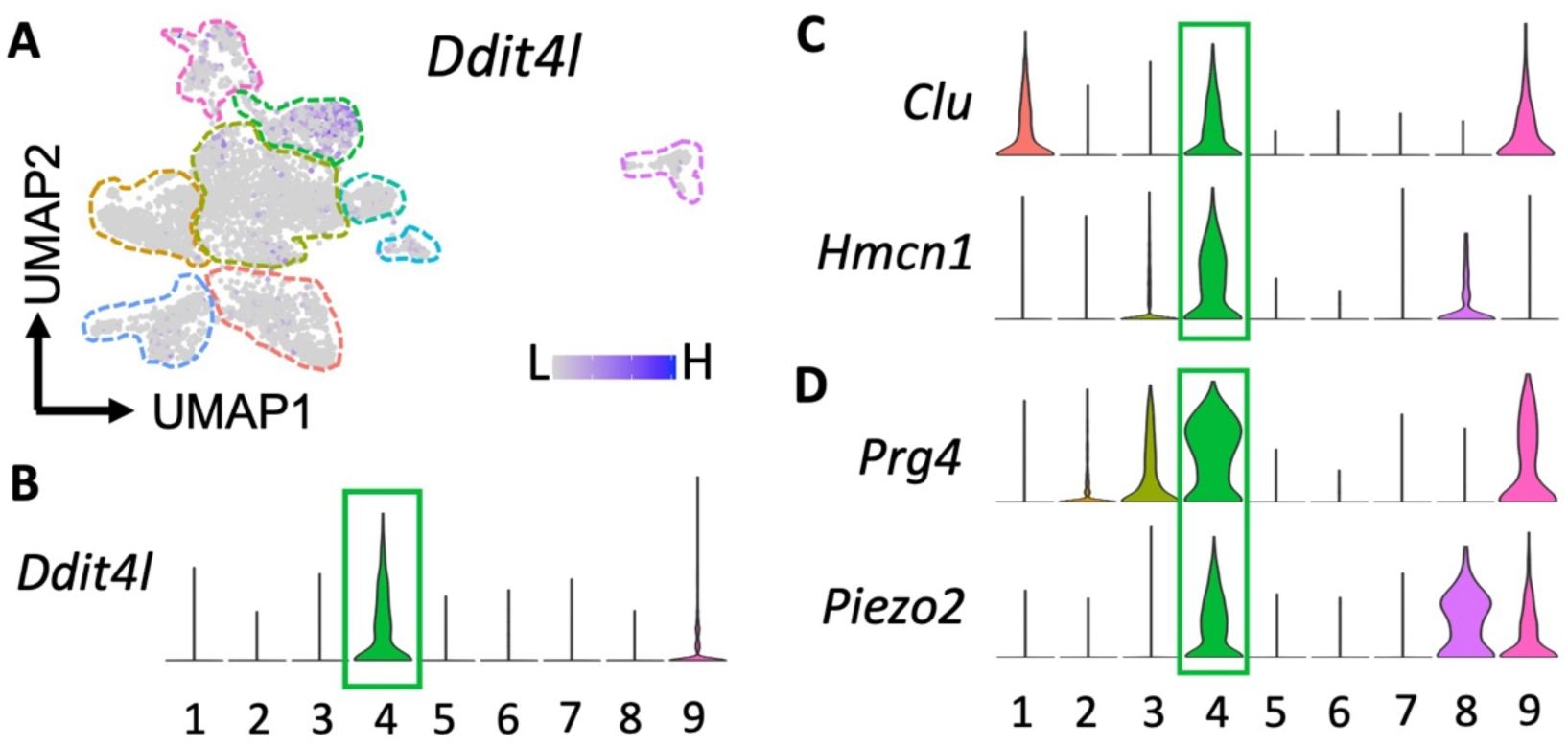
Cluster 4 mineralizing NMMCs are not involved in massive RCT pathologies. A) Feature plot on integrated UMAP and (B) violin plot for *Ddit4l.* B) Violin plots of previously defined markers of *Clu, Hmcn1* and (D) additional mineralizing markers *Prg4* and *Piezo2*.

**Supplemental Figure 5:**
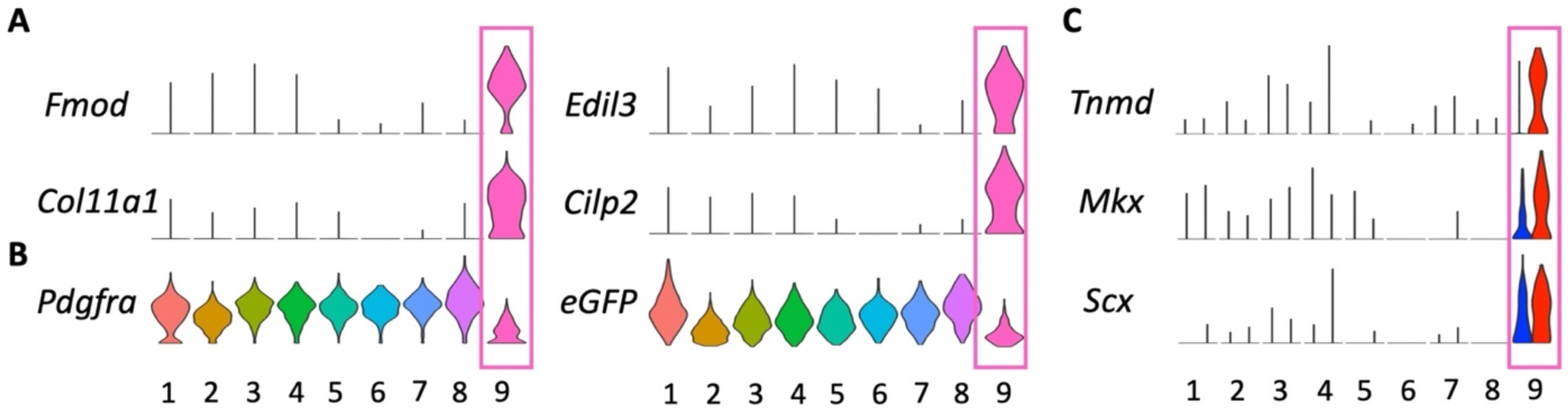
Cluster 9 tenocyte-like NMMCs are low *Pdgfra+* expressors and express more mature tenocyte markers after massive RCT. A) Violin plots for cluster 9 marker genes, *Fmod, Col11a1, Edil3,* and *Cilp2.* B) Violin plots for *Pdgfra* and *eGFP.* C) Split violin plots showing increases in tenocyte marker genes (*Tnmd, Mks,* and *Scx)* after massive RCT.

**Supplemental Figure 6:**
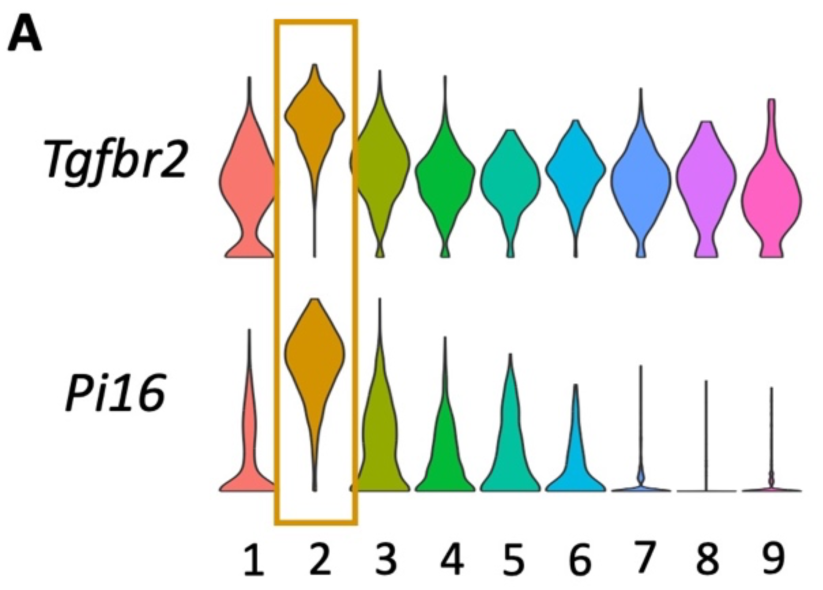
Additional published markers for *Dpp4+* cells. A) Violin plots of previously defined fibrotic markers for Dpp4+ NMMCs, *Tgfbr2* and *Pi16*

**Supplemental Figure 7:**
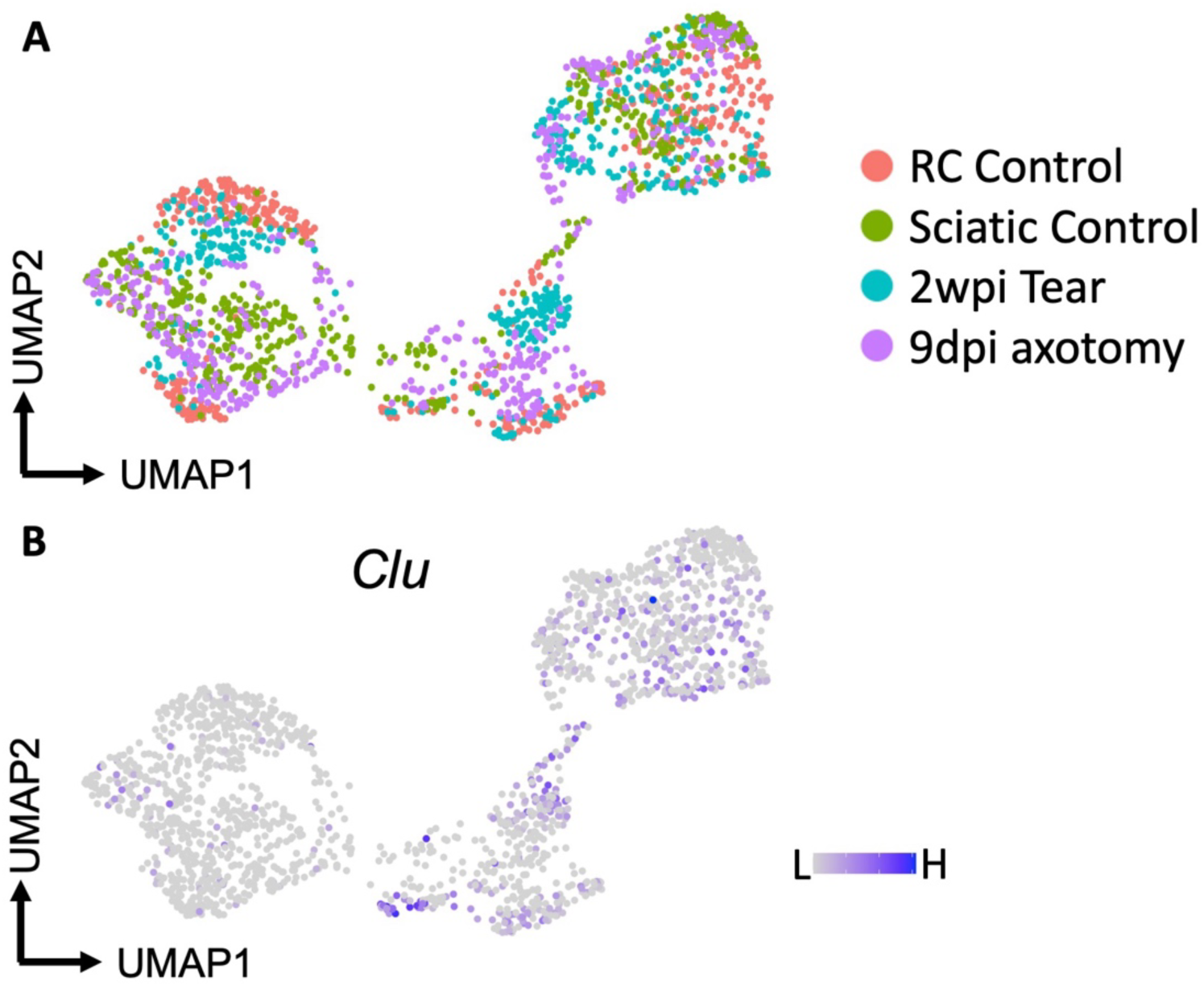
Nerve Associated NMMCs are likely PNS Nerve Fibroblasts. A) Overlaid data (RC uninjured control, 2wpi tear, Carr et al. uninjured, Carr et al. 9dpi axotomy) UMAP projection showing which dataset cells come from. B) Feature plot of *Clu* (cluster 4) cells intermingled with all data.

**Supplemental Figure 8:**
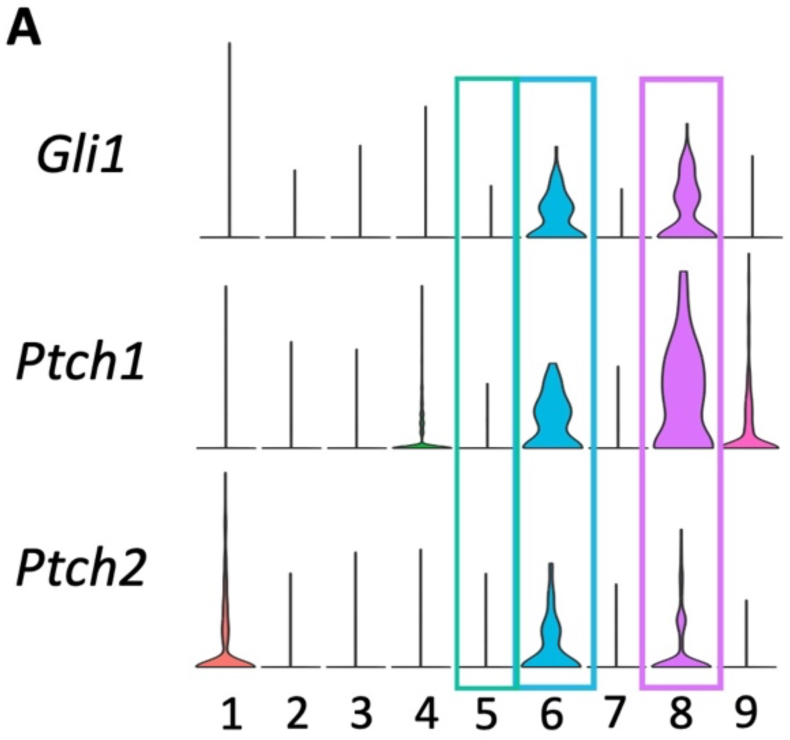
Nerve Associated NMMCs express Hh-responsive genes. A) Violin plots of Hedgehog Pathway genes, *Gli1, Ptch1,* and *Ptch2*

**Supplemental Figure 9:**
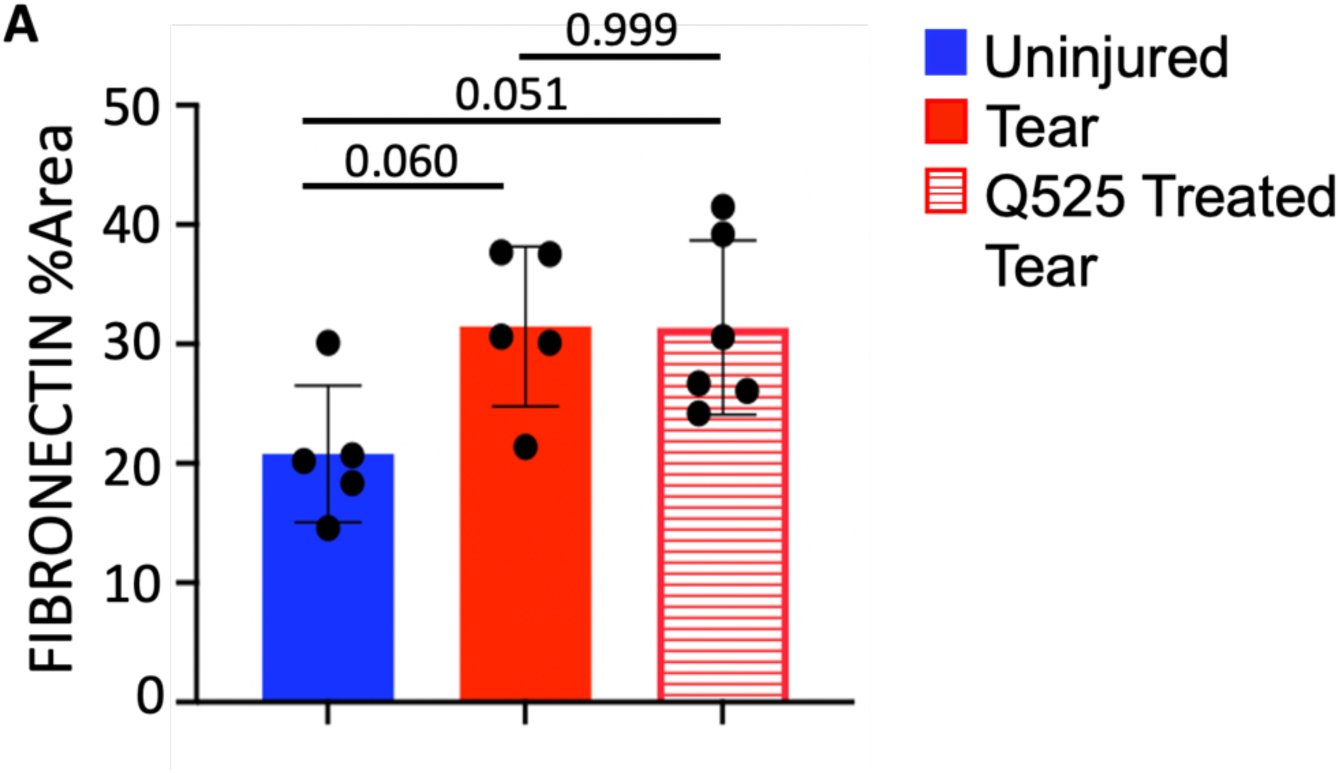
Treatment with RET agonist Q525 does not alter RCT induced fibrosis. A) Quantification of FIBRONECTIN staining for contralateral control, DMSO treated tear, and Q525 treated tear. Data are represented as mean±SD. n=5/6. Significance determined using one-way ANOVA with multiple comparisons. Control vs. DMSO tear p=0.060, Control vs. Q525 tear p=0.051, DMSO treated tear vs. Q525 treated tear p=0.999.

